# Morphological trait evolution in *Solanum* (Solanaceae): evolutionary lability of key taxonomic characters

**DOI:** 10.1101/2023.02.24.529849

**Authors:** Rebecca Hilgenhof, Edeline Gagnon, Sandra Knapp, Xavier Aubriot, Eric J. Tepe, Lynn Bohs, Leandro L. Giacomin, Yuri F. Gouvêa, Christopher T. Martine, Andrés Orejuela, Clara Inés Orozco, Iris E. Peralta, Tiina Särkinen

## Abstract

*Solanum* L. is one of the world’s largest and economically most important plant genera, including 1,245 currently accepted species and several major and minor crops (e.g., tomato, potato, brinjal eggplant, scarlet eggplant, Gboma eggplant, lulo, and pepino). Here we provide an overview of the evolution of 25 key morphological traits for the major and minor clades of this giant genus based on stochastic mapping using a well-sampled recently published phylogeny of *Solanum*. The most evolutionarily labile traits (showing >150 transitions across the genus) relate to plant structure (growth form and sympodial unit structure), herbivore defence (glandular trichomes), pollination (corolla shape and colour), and dispersal (fruit colour). Ten further traits show evolutionary lability with 50-100 transitions across the genus (e.g., specialised underground organs, trichome structure, leaf type, inflorescence position and branching, stamen heteromorphism). Our results reveal a number of highly convergent traits in *Solanum*, including tubers, rhizomes, simple leaves, yellow corollas, heteromorphic anthers, dioecy, and dry fruits, and some unexpected pathways of trait evolution that could be explored in future studies. We show that informally named clades of *Solanum* can be morphologically defined by trait combinations providing a tool for identification and enabling predictive phylogenetic placement of unsampled species.

## Introduction

*Solanum* L. is one of the world’s largest and economically most important plant genera with 1,245 currently accepted species (http://solanaceaesource.org/). Twenty-four of these are major crops (Knapp & al., 2004), including the potato (*S. tuberosum* L.), tomato (*S. lycopersicum* L.), and brinjal eggplant (*S. melongena* L.), as well as some lesser-known cultivated species, such as pepino (*S. muricatum* Aiton), lulo/naranjilla (*S. quitoense* Lam. and relatives), tree tomato or tamarillo (*S. betaceum* Cav.), cocona (*S. sessiliflorum* Dunal), scarlet eggplant (*S. aethiopicum* L.), Gboma eggplant (*S. macrocarpon* L.), and bush tomato (*S. centrale* J.M.Black).

Agriculture has benefited from the morphological diversity found in *Solanum* through exploitation of variation in traits such as underground storage organs and fruit morphology. Breeding programmes have also used the wide range of diversity present in gene pools of the various crops and crop wild relatives (CWRs), to enhance quality and yield (e.g., Gur & Zamir, 2004; Semel & al., 2006; Lippman & Zamir, 2007), abiotic stress tolerance, and disease resistance in cultivated potato, tomato, and eggplant (e.g., Prohens & al., 2013, 2017; Dempewolf & al., 2017; Villanueva & al., 2021). One of the best examples of the use of wild diversity in crop breeding is the single cross between the cultivated tomato and *S. habrochaites* D.M.Spooner & S.Knapp, a green-fruited wild tomato relative from the northern Andes, that increased fruit soluble solid content in tomato by 22% and brought significant profit for industry (Tanksley & al., 1996; Tanksley & McCouch, 1997; Bernacchi & al., 1998).

*Solanum* has served as a model system for research into the genetic basis of several important morphological traits. Examples include quantitative trait locus (QTL) mapping studies of major crop species, which have helped to explore morphological traits relevant to plant breeding (e.g., D’Hoop & al., 2008, 2014), as well as evolutionary developmental studies involving traits such as genetic control of fleshy fruits (Pabón-Mora & Litt, 2011; Tomato Genome Consortium, 2012), anther cone cohesion (Glover & al., 2004), leaf shape and lobing (e.g., Geeta & al., 2011; Chitwood & al., 2013; Wu & al., 2018; Nakayama & al., 2021), breeding systems (self-incompatibility and clonality; Vallejo-Marín & O’Brien, 2007), as well as traits related to chemical defences and animal-plant interactions (e.g., Tingey & Gibson, 1978; Tingey & Laubengayer, 1981; Avé & Tingey, 1986; D’Hoop & al., 2008), and pathogen resistance (e.g., Gebhardt & Valkonen, 2001; Jupe & al., 2012; Thaler & al., 2012).

Despite agricultural interest and ongoing research, much of the morphological diversity in *Solanum* (Fig. 1) remains underutilized and unexplored. Species of *Solanum* are highly variable in both vegetative and reproductive morphology, including their growth form which ranges from ephemerals in the world’s driest deserts to tiny annual herbs growing at 4,000 m elevation to large trees and herbaceous climbers from premontane and lowland rainforests (Fig. 1A-F). New and unexpected morphological diversity continues to be discovered as baseline taxonomic work advances in *Solanum*, with some of recent discoveries including a species with heart-shaped anthers (*S. anomalostemon* S.Knapp & M.Nee; Knapp & Nee, 2009), tuber-bearing shrubs (Asterophorum clade; Gouvêa & Stehmann, 2019), species with leaky dioecy and fluid sex expression (Martine & al., 2009; McDonnell & al., 2019) and with resin-glands (Silva Sampaio & al., 2021).

**Fig. 1.**
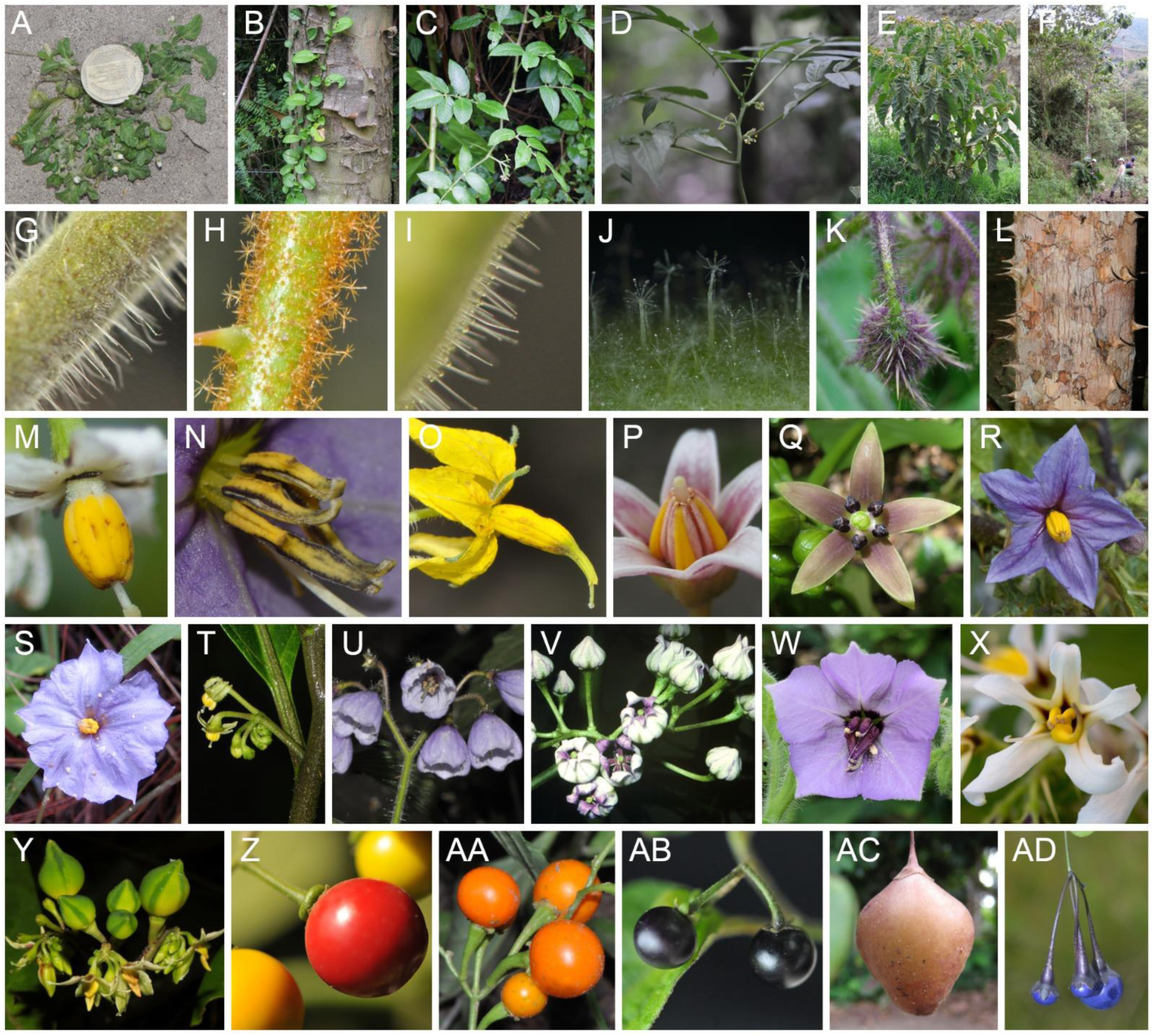
Morphological diversity across *Solanum*. **A**, Annual herb (Morelloid clade, *S. weddellii* Phil.); **B**, Herbaceous vine (Herpystichum clade, *S. brevifolium* Dunal; **C**, Woody vine (Tomato clade, *S. juglandifolium* Dunal); **D**, Single-stemmed shrub (Pteroidea clade, *S. mite* Ruiz & Pav.); **E**, Shrub (Torva clade, *S. glutinosum* Dunal); **F**, Tree (Crinitum clade, *S. S. sycophanta* Dunal); **G**, Simple glandular trichomes (Tomato clade, *S. habrochaites* S.Knapp & D.M.Spooner); **H**, Stellate glandular trichomes (Torva clade, *S. asperolanatum* Ruiz & Pav.); **I**, Mix of simple glandular (short) and eglandular (long) trichomes (Tomato clade, *S. arcanum* Peralta); **J**, Stellate glandular trichomes (Erythrotrichum clade, *S. kollastrum* Gouvêa & Giacomin); **K**, Needle-like prickles on calyx (EHS clade, *S. dasyphyllum* Schumach. & Thonn.); **L**, Broad-based prickles on trunk (Crinitum clade, *S. kioniotrichum* Bitter ex J.F.Macbr.); **M**, Monomorphic stamens, most common state in *Solanum* (Anarrhichomenum clade, *S. appendiculatum* Dunal); **N**, Apical and basal anther modifications (i.e., horn-like projections; Normania clade, *S. trisectum* Dunal); **O**, Apical anther modifications (i.e., appendages; Tomato clade, *S. corneliomulleri* J.F.Macbr.); **P**, Enlarged anther connectives (Pachyphylla clade, *S. betaceum* Cav.); **Q**, Deeply stellate purple corollas (Pachyphylla clade, *S. sycocarpum* Mart. & Sendtn.); **R**, Broadly stellate purple corollas (EHS clade, *S. linnaeanum* Hepper & P.-M.L. Jaeger); **S**, Rotate purple corollas with abundant interpetalar tissue (Herpystichum clade, *S. trifolium* Dunal); **T**, Deeply stellate yellow-green corollas lacking interpetalar tissue (Pteroidea clade, *S. anceps* Ruiz & Pav.); **U**, Campanulate pale lilac corollas (Morelloid clade; *S. fiebrigii* Bitter); **V**, Urceolate white-purple corollas (Pachyphylla clade, *S. diversifolium* Dunal); **W**, Bilaterally symmetric corollas with heteromorphic anthers (Normania clade, *S*. *trisectum* Dunal); **X**, Bilaterally symmetric corollas with heteromorphic anthers (Androceras clade; *S. grayi* Rose var. *grandiflorum* Whalen); **Y**, Obovoid, apically pointed fleshy berries (Thelopodium clade, *S. thelopodium* Sendtn.); **Z**, Globose fleshy berries with colour variation through maturation from yellow (unripe) to red (fully mature; Cyphomandropsis clade, *S. amotapense* Svenson); **AA**, Globose orange berries (Reductum clade, *S. reductum* C.V.Morton); **AB**, Globose black berries (Morelloid clade, *S. longifilamentum* Särkinen & P.Gonzáles); **AC**, Obovoid, apically pointed brown berries (Herpystichum clade, *S. limoncochaense* Tepe); **AD**, Globose blue berries (Dulcamaroid clade, *S. laxum* Spreng.). Photo vouchers: **A** *Särkinen 4038*, **B** *Tepe 2726*, **C** *Farruggia 3998*, **D** *Särkinen 4822*, **E** *Knapp 10594*, **F** *Tepe 2327,* **G** *Särkinen 4524,* **H** *Knapp 10336,* **I** *Särkinen 4503,* **J** *Gouvêa 280,* **K** *Särkinen 4536,* **L** *Melchor Castro 1446,* **M** *Knapp 10156*, **N** *Nijmegen 984750158*, **O** *Knapp 10212*, **P** *Tepe s.n.*, **Q** *Bohs s.n.*, **R** Knapp s.n., **S** *Tepe 2684*, **T** *Fajardo et al. 3982*, **U** *Barboza et al. 3548*, **V** *Benitez de Rojas 2744*, **W** *cult. Madeira*, **X** *Vallejo-Marin 08-s-78,* **Y** *Melchor Castro 1454*, **Z** *Särkinen 4508*, **AA** *Barboza 3516*, **AB** *Särkinen s.n.*, **AC** *Tepe 2627,* **AD** *Giacomin 1737*. Photographs by S. Knapp, T. Särkinen, P. Gonzáles Arce, E. Tepe, L. Bohs, Y.F. Gouvêa, M. Vallejo-Marin, M. Benedito, L. Giacomin, and A. Fuentes.

*Solanum* has been traditionally characterised by its relatively uniform floral morphology with sympetalous, five-parted flowers with a central anther cone of poricidally dehiscent anthers (Fig. 1M-R). Prior to molecular phylogenetic studies, the genus was divided into sections based on morphological characters (Dunal, 1852; D’Arcy, 1972; Hunziker, 2001); most of these divisions have subsequently been shown to be para- or polyphyletic (Olmstead & Palmer, 1997; Bohs, 2005; Weese & Bohs, 2007). These traditional formal infrageneric systems have since been replaced by a system of informally named clades that reflect monophyletic groups (Fig. 2; Bohs, 2005; followed by subsequent studies, e.g., Stern & al., 2011; Särkinen & al., 2013; Tepe & al., 2016; Gagnon & al., 2022). Morphological characterisation of some of these clades has been difficult, however, and the robustness of proposed morphological synapomorphies has not been tested. The use of DNA sequence data has also changed the circumscription of *Solanum* by showing that taxa with stamen heteromorphism and/or anther modifications previously segregated for these characters are nested within the genus (e.g., *Lycopersicon* Mill., *Normania* Lowe, *Cyphomandra* Mart. ex Sendtn.; Spooner & al., 1993; Bohs & Olmstead, 1997; Olmstead & Palmer, 1997; Bohs & Olmstead, 2001; Tepe & al., 2016). These changes have had a minimal effect on the size of *Solanum* but have expanded the morphological diversity included within the genus, especially in relation to androecium characteristics.

**Fig. 2.**
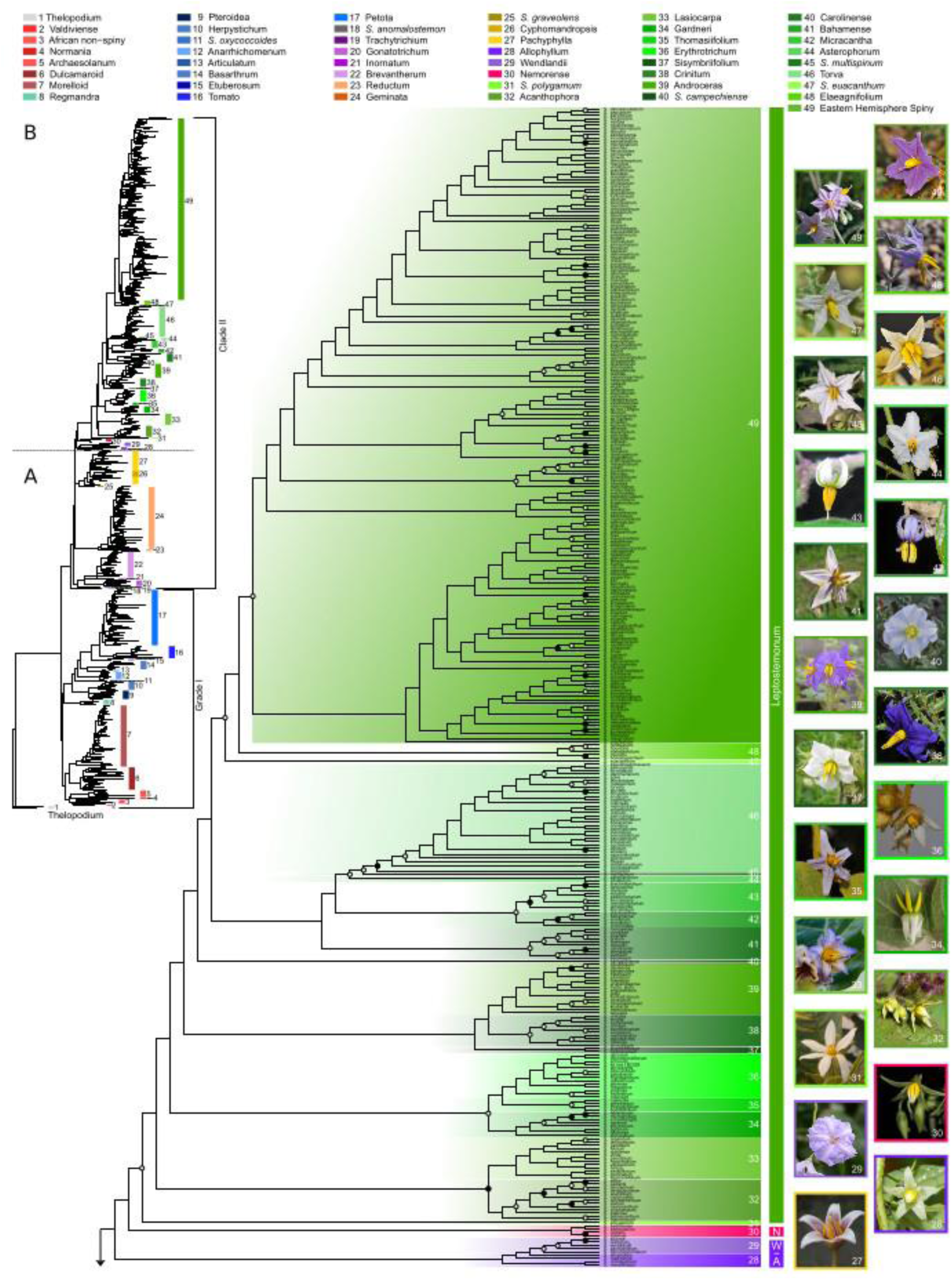
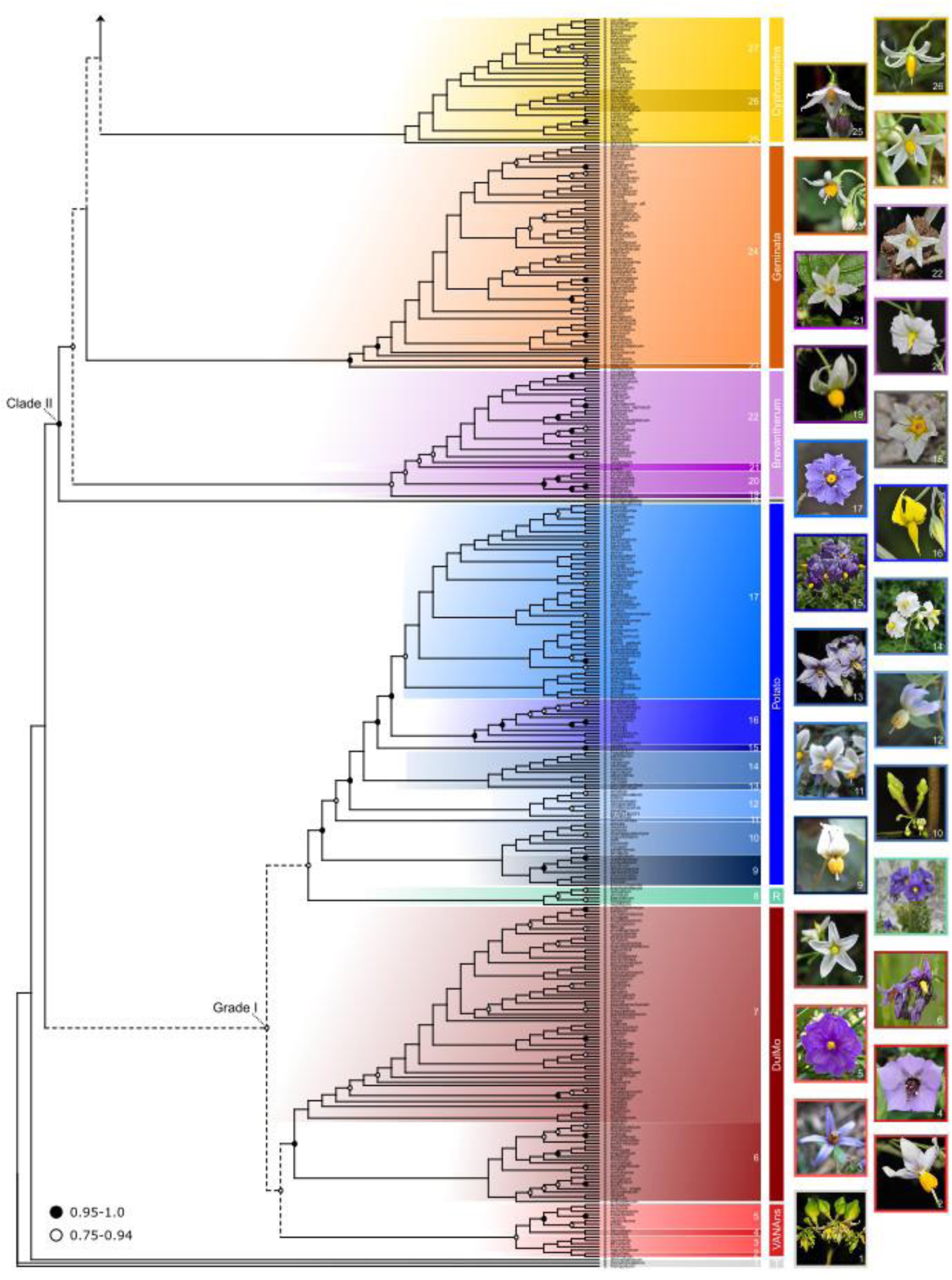
Phylogeny of *Solanum* highlighting the informally named infrageneric clades based on Maximum Likelihood analysis (RaxML) of 742 *Solanum* species (60% of total known diversity) with two nuclear and seven plastid regions by Gagnon & al. (2022). Infrageneric clades are colour-coded and numbered reflecting the currently recognised major and minor clades of *Solanum* (Table 1): bright red shades highlight minor clades within VANAns clade, dark reds DulMo, blues Potato clade, purples Brevantherum (subfigure A), orange shades Geminata, yellows Cyphomandra, purple Wendlandii-Allophyllum (subfigure B), pink Nemorense, and green shades indicate minor clades within the large Leptostemonum clade. Nodes without circles have maximum branch support (100% boostrap), nodes with black circles strong support (>95%), and nodes with white circles moderate to weak support (75–94%). Dashed lines indicate nodes with nuclear-plastome discordance highlighted in Gagnon & al. (2022) collapsed in our analyses (Figs. 4-7).

**Table 1.** Results of the ancestral trait reconstruction analyses (stochastic mapping) of 25 morphological traits across Solanum. Mean number of changes modelled across 200 simulations is given based on the best model of the three models tested identified using the Akaike Information Criteria (AIC): Equal Rates (ER), Symmetric (SYM), and All Rates Different (ARD). Traits were categorised as evolutionarily highly labile (>150 transitions; darkest grey), labile (50-100 transition; light grey), conserved (10-50 transitions), or highly conserved (<10 transitions). The 95% highest probability density (HPD) of the mean number of transitions is shown for each trait. Details of the estimated transition rates of the underlying Markov models of each trait can be found in SI Table 3.

|  | Trait | States | Best model(s) based on AIC | Observed mean number of transitions (95% HPD) | Results | Main references used for scoring states |
| --- | --- | --- | --- | --- | --- | --- |
| 1. | Growth form | Herb, shrub, single-stemmed shrub, tree, herbaceous vine, woody vine, epiphyte | ARD | 109.9 (xx) | Highly evolutionarily labile with >100 transitions |  |
| 2. | Specialised underground organs | Rhizomes, underground storage organs, absent | ARD | 64.4 (xx) | Evolutionarily labile with 50-100 transitions | Spooner & al., 2004, 2016, 2019; Bennett, 2008; Knapp & al., 2017 |
| 3. | Prickles | Broad-based, needle-like, absent | ARD | 82.2 (xx) | Evolutionarily labile with 50-100 transitions | Seithe, 1962, 1979 |
| 4. | Trichome structure | Simple, dendritic, stellate, absent | ARD | 93.8 (xx) | Evolutionarily labile with 50-100 transitions | Seithe, 1962, 1979; Roe, 1971; Seithe & Anderson, 1982; Freire de Carvalho & |
|  |  |  |  |  |  | Machado, 1991; Giacomini & Stehmann, 2011; Stern & Bohs, 2012; Watts & Kariyat, 2022 |
| 5. | Glandular trichome type | Simple glandular, stellate glandular, absent | ARD | 208.3 (xx) | Highly evolutionarily labile with >100 transitions | Peralta & al., 2008; Kang & al., 2014; Watts & Kariyat, 2022 |
| 6. | Pseudostipules | Present (pair), present (single), absent | SYM | 9.2 (xx) | Highly conserved with <10 transitions | Spooner & al., 2004, 2016, 2019; Peralta & al., 2008 |
| 7. | Sympodial unit structure | Plurifoliate, trifoliate, difoliate non-geminate, difoliate geminate, unifoliate | ARD | 215.0 (xx) | Highly evolutionarily labile with >100 transitions | Danert, 1958; Child, 1979 |
| 8. | Leaf division (i.e., type) | Simple, lobed, compound | ARD | 73.4 (xx) | Evolutionarily labile with 50-100 transitions | Geeta & al., 2011 |
| 9. | Inflorescence position | Terminal, internodal, axillary, leaf-opposed, spur-shoots | SYM | 93.6 (xx) | Evolutionarily labile with 50-100 transitions | Danert, 1958; Child, 1979 |
| 10. | Inflorescence branching | Unbranched, forked, multi-branched | ARD | 89.4 (xx) | Evolutionarily labile with 50-100 transitions | Lippman & al., 2008 |
| 11. | Sexual system | Cosexual (hermaphroditic), andromonoecious, dioecious | ARD | 87.6 (xx) | Evolutionarily labile with 50-100 transitions | Symon, 1970, 1979a,b; Levine & Anderson, 1986; Whalen & Costich, 1986; Anderson & Symon, 1989; Knapp & al., 1998; Dupont & Olesen, 2006; Anderson & al., 2015; Ndem-Galbert & al., 2021 |
| 12. | Corolla shape | Deeply stellate, broadly stellate, rotate | SYM | 272.0 (xx) | Highly evolutionarily labile with >100 transitions |  |
| 13. | Corolla bilateral symmetry | Present, absent | ARD | 15.1 (xx) | Conserved with 10-50 transitions | Bohs & al., 2007 |
| 14. | Corolla colour | White, green, purple (including blues and pinks), yellow (including yellowish) | ARD | 180.2 (xx) | Highly evolutionarily labile with >100 transitions | Passarelli & Bruzzone, 2004 |
| 15. | Stamen heteromorphism | Filaments, anthers, absent | ARD | 50.1 (xx) | Evolutionarily labile with 50-100 transitions | Bohs & al., 2007 |
| 16. | Anther connectives | Enlarged, not enlarged | ARD | 7.9 (xx) | Highly conserved with <10 transitions | D'Arcy & al., 1990; Bohs, 1994; Cocucci, 1996 |
| 17. | Anther modifications | Appendage, beak, horn, sack, absent | ER | 7.1 (xx) | Highly conserved with <10 transitions | Francisco-Ortega & al., 1993; Peralta & al., 2008; Knapp & Nee, 2009; Barboza, 2013 |
| 18. | Anther shape | Cylindrical, tapered, cordate | SYM | 12.2 (xx) | Conserved with 10-50 transitions |  |
| 19. | Pedicel insertion | Flat, cup-shaped | ER | 2.0 (xx) | Highly conserved with <10 transitions |  |
| 20. | Pedicel articulation | Basal, near-basal, basal 1/4-1/2, distal 1/2, absent | ER | 10.3 (xx) | Conserved with 10-50 transitions |  |
| 21. | Fruit type | Fleshy berry, dry berry | ER | 13.5 (xx) | Conserved with 10-50 transitions | Symon, 1979a, 1984; Whalen, 1984; Knapp, 2002b; Chiarini & Barboza, 2007 |
| 22. | Fruit colour | Green, white, purple, yellow, orange, red | ARD | 284.0 (xx) | Highly evolutionarily labile with >100 transitions | Peralta & al. 2008; Dhar & al., 2015 |
| 23. | Fruit trichomes | Present, absent | ARD | 67.0 (xx) | Evolutionarily labile with 50-100 transitions |  |
| 24. | Fruiting calyx modifications | Inflated, appressed, swollen, absent | ARD | 61.9 (xx) | Evolutionarily labile with 50-100 transitions |  |
| 25. | Stone cells | Present, absent | ARD | 42.2 (xx) | Conserved with 10-50 transitions | Bitter, 1911, 1914; Symon, 1994; Knapp, 2002b; Särkinen & al., 2018; Knapp & al., 2019 |

The current clade-based informal infrageneric system divides *Solanum* into 49 lineages (called here minor clades), which are grouped into 12 larger clades (called here major clades) and further into three main groups (Figs. 2 & 3; Bohs, 2005; Gagnon & al., 2022): (1) the small Thelopodium clade that consists of three species sister to the rest of *Solanum*; (2) Grade I, previously referred to as Clade I (Särkinen & al., 2013) with ca. 339 non-spiny species including the cultivated tomato, potato and pepino; and (3) Clade II, the largest monophyletic lineage in the genus that includes 73% of *Solanum* (903 currently accepted species) including the tree tomatoes and all cultivated eggplants and their relatives. Within Grade I are 4 major and 16 minor clades (Fig. 2); major clades are Regmandra, VANAns (Valdiviense, Archaesolanum, Normania, and African non-spiny), DulMo (Dulcamaroid and Morelloid), and the Potato clade that includes several economically important minor clades (e.g., Tomato, Petota, Etuberosum, Basarthrum; Gagnon & al., 2022). Clade II contains seven major clades, including the large Leptostemonum clade with ca. 580 species, and 32 minor clades including the Eastern Hemisphere Spiny clade (hereafter EHS, previously known as the Old World clade; Fig. 2; Gagnon & al., 2022).

**Fig. 3.**
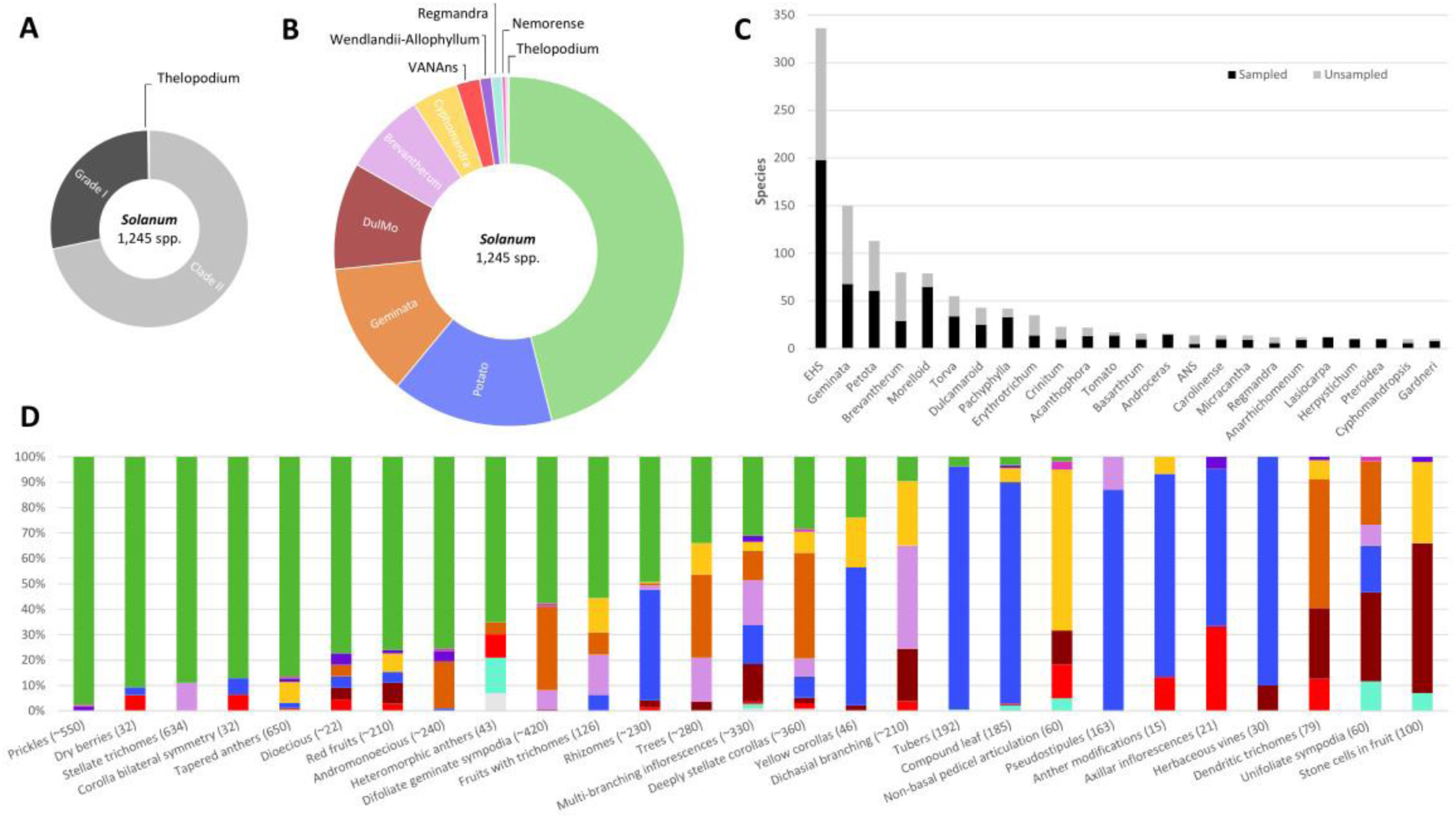
Overview of morphological diversity across *Solanum* and the informally named infrageneric groups referred to as major and minor clades. **A**, Division of *Solanum* into three main lineages and an overview of species diversity within these large groups; **B**, Species diversity across major clades of *Solanum*; **C**, Sampling density across the largest minor clades of *Solanum* in the current species-level molecular phylogeny (Gagnon & al., 2022) showing clades with **≥**10 currently accepted species. Total number of accepted species is reported for each clade based on current taxonomy in SolanaceaeSource; **D**, Trait diversity across major clades of *Solanum*, showing the estimated proportion of species from each major clade that possess the traits visualised including species not yet sampled in the phylogeny. Clade colouring in B and D follows Figure 2. EHS stands for Eastern Hemisphere Spiny clade previously known as Old World Spiny.

### Introduction to general morphology of *Solanum* here

Here we examine morphological trait evolution across *Solanum* clades with the aim of: (1) providing an overview of current morphological knowledge and diversity across the genus in a phylogenetic context based on the well-sampled phylogeny of Gagnon & al. (2022); (2) identifying evolutionarily labile and conserved traits across *Solanum*; and (3) providing morphological descriptions for all *Solanum* clades. Our study is inspired by the dynamic research community working on *Solanum* continuously generating new knowledge on morphological and genotypic diversity as well as unravelling the molecular mechanisms that control the genotype-phenotype interactions across the genus. More than 100 new species have been described in *Solanum* over the past 10 years (e.g., Tepe & al., 2012; Bean, 2014; Giacomin & Stehmann, 2014; Stern & al., 2014; Knapp & al., 2015; Särkinen & al., 2015a, b, c; Bean, 2016a, b; Martine & al., 2016; Särkinen & Knapp, 2016; Knapp & Särkinen, 2018; Gouvêa & Stehmann, 2019; Knapp & al., 2020; Aubriot & Knapp, 2022), revealing new traits and trait combinations. This, coupled with the densely sampled molecular phylogeny, provides an ideal opportunity to study the origin and evolution of major vegetative and reproductive traits in this agronomically important genus, and to assess/confirm diagnostic traits for *Solanum* clades using a formal analysis with standardised terminology. Results are discussed in relation to previous evolutionary and developmental hypotheses, and new morphological hypotheses of trait evolution and homology are proposed that could be tested with further studies and new tools.

## Materials and Methods

### Supermatrix topology

The phylogenetic framework is based on the combined supermatrix assembled by Gagnon & al. (2022) that includes 742 species of *Solanum* (60% of the 1,245 currently accepted species; SI Table 1) and covers all minor clades with 35-100% species sampling in each (Fig. 3A-C, SI Table 1; Gagnon & al., 2022). All outgroup sequences of *Jaltomata* Schltdl. were excluded, because coding all morphological diversity present in the sister genus was beyond the scope of this study. Instead, we incorporated the uncertainty of the root state in our ancestral state reconstruction methods (see below). Nodes corresponding to areas of high nuclear-plastome discordance along the backbone of Grade I and Clade II in phylogenomic analyses (Gagnon & al., 2022) were collapsed using the TreeTools package (Smith, 2019) to account for phylogenetic uncertainty and topological conflict.

### Morphological traits

A total of 25 morphological traits was evaluated, including eight vegetative traits, sexual system (sensu Cardoso & al., 2018), two inflorescence traits, seven floral traits, and seven fruit-related traits (Table 1; SI Table 2). We selected the traits for scoring because they have been previously used in *Solanum* taxonomy and are of potential interest to the wider research community and in future crop breeding programs. All species present in the phylogeny (after excluding 17 unpublished species for which full morphological descriptions are not yet available) resulted in 725 coded *Solanum* species (58% of all species; Fig. 3D, SI Table 2). Scoring was done from the taxonomic literature and herbarium specimens included in the SolanaceaeSource database (PBI Solanum Project, 2022) identified by *Solanum* experts (SI Table 1; e.g., Symon, 1994; Knapp & Helgason, 1997; Contreras & Spooner, 1999; Bennett, 2008; Peralta & al., 2008; Tepe & Bohs, 2011; Knapp, 2013; Knapp & Vorontsova, 2016; Särkinen & al., 2018; Knapp & al., 2019). Terminology from descriptions was standardized based on our (sometimes incomplete) knowledge of character homology. Polymorphisms were included in all traits when multiple states were known to be present within a species.

Growth in *Solanum* is sympodial, with a terminal inflorescence followed by continued shoot growth from axillary buds (Danert, 1958, 1970) often giving the stems a zig-zag appearance, where leaves can be paired at nodes (geminate). We coded sympodial structure following Danert (1970) with numbers of leaves (1, 2, 3 or many) between each inflorescence, coupled with whether leaves are found in pairs along each node (i.e., geminate or not; Table 1). Inflorescence position is determined by variation in fusion of inflorescence and stem axes (Danert, 1958); internodal inflorescences were coded as those that arise along the stem not associated with leaves (axillary buds), while leaf-opposed inflorescences arise in conjunction with a leaf or leaf pair.

Trichome structure was coded combining the developmental pathways suggested by Seithe (1962, 1979) and the structure-based terminology of Roe (1971). Simple trichomes included all unbranched trichomes including 2-celled “bayonet” trichomes (Fig. 1G; Seithe & Anderson 1982). All branched or forked trichomes and echinoid dendritic trichomes (e.g., Knapp, 2002a) were scored as dendritic. Stellate trichomes included geniculate trichomes (simple trichomes derived from stellate trichomes), bristles, densely compacted multi-angulate stellate trichomes that have been referred to as echinoid, as well as lepidote scales developmentally derived from stellate trichomes (Fig. 1H & J; Seithe, 1962, 1979; Stern & al., 2013). Presence of glandular tips on trichomes was coded separately because they are found on all trichome types independent of structure (Fig. 1I-J). The degree of leaf lobing was categorised by roughly quantifying the depth of the sinus to the distance to the midrib. Simple leaves included leaf blades with entire, serrate, or crenate margins. Leaf blades lobed a quarter to halfway to the midrib were coded as lobed. Compound leaves were defined as leaves with blades divided all the way to the midrib, although leaflets in most of these *Solanum* species are variously decurrent along the leaf midrib thus leaves are more strictly pinnatifid.

*Solanum* species have a wide variety of sexual (breeding) systems (sensu Cardoso & al., 2018), species recorded to have both short- and long-styled flowers on the same plant were coded as andromonoecious, and species with individuals possessing only short- or long-styled flowers were coded as dioecious; all other species were coded as cosexual (i.e., hermaphroditic). Three states of corolla shape were recognised reflecting the amount of interpetalar tissue between lobes: deeply stellate corollas without apparent interpetalar tissue (i.e., deeply stellate, Fig. 1P-Q), and two states with increasing amount of interpetalar tissue, broadly stellate (Fig. 1R) and rotate (Fig. 1S & W, U), respectively. Our coding does not take into account the orientation of corolla lobes as corolla lobes in *Solanum* often change orientation through anthesis but is a measure of corolla division only. For example, the urceolate corollas of *S. diversifolium* Dunal were coded as deeply stellate based on the lobe length relative to the tube (Fig. 1V). Corolla colour was coded as four states: white, green, purple (including blues and pinks), and yellow (including pale, dull and bright yellows). The colour of the central “eye” of the corolla was not included in the coding, and corollas described with multiple colours in species descriptions (e.g., yellow-green) were coded as multistate (yellow + green). We recognised four different anther modifications for coding purposes, homology of these states has not been tested yet: long basal anther projections referred to as horns, short basal anther extensions referred to as sacks, long fused (i.e., connivent) apical appendages, and short loose (i.e., not connivent) apical anther beaks.

Fruit type was coded as either fleshy (including juicy, spongy, or woody indehiscent berries as well as the explosively dehiscent berries of the Gonatotrichum clade that contain fleshy mesocarp when fully ripe) or dry (including all dehiscent berries that lack fleshy mesocarp when fully ripe). Fruiting pedicel articulation was coded as three states based on distance from the inflorescence rhachis (SI Table 1). Fruiting calyx modifications were coded as four states: inflated (papery, lantern- or balloon-like enlarged calyces loosely covering at least >50% of the fruits), appressed (enlarged calyces tightly surrounding at least 50% of the fruits), swollen (thickened calyx tube and/or lobes often with a doughnut shaped ring of swelling; Fig. 1Z), or absent. Mature fruit colour was coded based on the external peel colour (i.e., not accounting for flesh colour) with six states following corolla colour with the addition of orange and red (Fig. 1Y-AD). Species with striped or multi-coloured mature berries, as well as those that change colour more than once (e.g., immature fruits green, intermediately mature fruits yellow, and mature fruits red) were scored as polymorphic. Presence of stone cells in fruits was coded from taxonomic monographs, field observations, or herbarium specimens similar to all other characters, as they can be easily seen (and felt) in both fresh and dried specimens if they are present.

### Trait evolution

Stochastic character mapping (SIMMAP; Huelsenbeck & al., 2003; Bollback, 2006) was performed using phytools v.0.6-99 package (Revell, 2012) in R (R Core Team, 2021). All traits were treated as unordered and equally weighted. Polymorphic clades were coded with equal probability of all states found to be present. Three different transition rate models were used: (1) Equal Rates (ER), i.e., a single rate for all possible transitions between states; (2) Symmetrical Rates (SYM); and (3) All Rates Different (ARD). For binary characters, only ER and ARD were run. The best fitting model was identified using the Akaike Information Criterion (AIC; Akaike, 1974) in phytools. The root state was treated as equally likely for all characters (pi=equal). Stochastic mapping was simulated with 200 generations to obtain a posterior probability distribution of ancestral states across the tree. The mean number of transitions per trait was determined based on the best fitting model. We interpreted those traits with >50 transitions as labile, and those with <50 transitions were interpreted as conservative.

## Results

Six traits show more than 150 mean shifts across *Solanum* indicating high evolutionary lability. These relate to plant structure (growth form and sympodial unit structure), herbivore defence (glandular trichomes), pollination (corolla shape and colour), and dispersal (fruit colour; Table 1; Fig. 4; SI Table 3). Switches between herbaceous and woody growth forms (shrubs, woody vines, and trees) are frequent and most common, with one origin of epiphytism from herbaceous ancestors (Fig. 4A). Changes in sympodial unit structure are frequent, relating to changes in growth form by enabling the modification of growth unit size, with difoliate geminate structure being modelled as ancestral in *Solanum* (Fig. 4B). Plurifoliate sympodial units dominate in Grade I and difoliate (geminate and non-geminate) units are more frequent in Clade II (Fig. 4B). Presence of glandular trichomes is highly labile across *Solanum*, with multiple independent origins and losses of both simple and stellate glandular trichomes (Table 1; SI Table 3; Fig. 4C). Corolla shape shows >270 transitions across *Solanum* with changes driven by the amount of interpetalar tissue present (Fig. 4D). Colour in both corollas and fruits shows high evolutionary lability, with most colours convergent in both corollas and fruits (Fig. 4E-F). Purple corollas are modelled to be ancestral in most clades in Grade I, while in Clade II white corollas are as ancestral (Fig. 4E). Loss and gains of purple pigments are the most common, while transitions to yellow are less common with a minimum of five independent origins in *Solanum* from both white and purple ancestors (Fig. 4E; SI Table 3). Fruit colour shows 284 transitions across *Solanum* with frequent shifts from yellow to green, yellow to red, and purple to green, and a minimum of 58 independent origins of red fruits (Fig. 4F; SI Table 3).

**Fig. 4.**
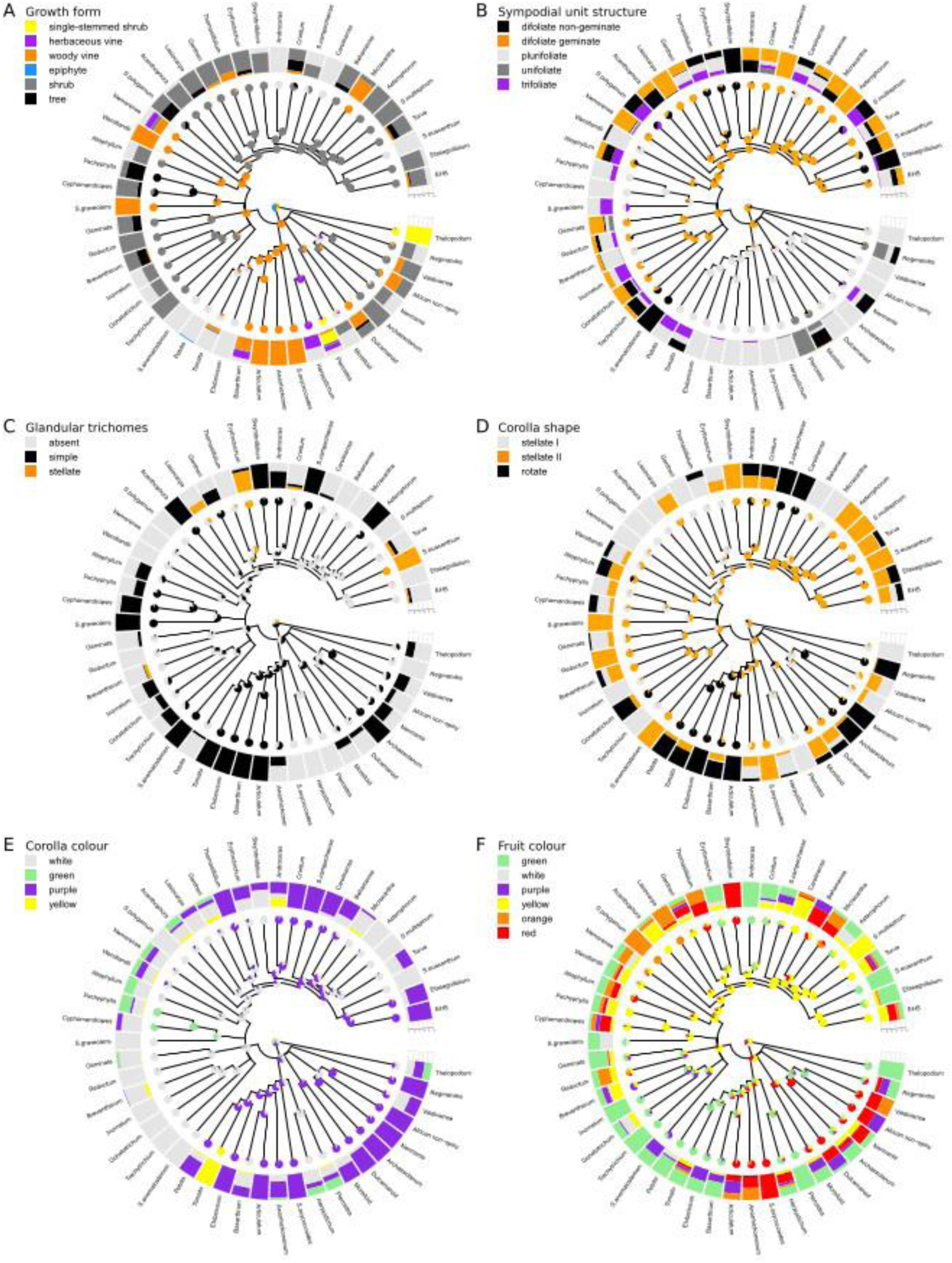
Evolution of the most highly labile morphological traits in *Solanum* based with >150 transitions species-level analysis using stochastic character mapping. **A**, Growth form; **B**, Sympodial unit structure; **C**, Glandular trichomes; **D**, Corolla shape; **E**, Corolla colour; and **F**, Fruit colour. Results from the best model are shown for each character (see Table 2 for details) based on 200 simulations. The topology used for mapping was derived from a supermatrix phylogeny with nine loci (two nuclear and seven plastid loci; Gagnon & al., 2022) with 725 species sampled and coded for each trait (60% of all species). All major and minor clades are labelled; tips reflect the crown nodes of each minor clade. Piecharts indicate likelihood of modelled ancestral states along the nodes, and frequency bars (tips) reflect proportion of species sampled within each clade with each state.

**Table 2.** Definition/description of Solanum clades based on morphology, geography, and broad-scale ecology. In parentheses after each clade name is the current number of species postulated to be in each. Numbers of species in each clade sampled in our phylogeny can be found in SI Table 1.

| Main group | Description | Major clade | Description | Minor clade | Description | Geography |
| --- | --- | --- | --- | --- | --- | --- |
| **Thelopodium (3)** |  |  |  |  | Single-stemmed (unbranched) shrubs with heteromorphic anthers and filaments and beak-like anther modifications, fruits apically pointed onion dome shaped and pointing upwards | lowland and pre-montane South America |
| **Grade I** | Non-prickly herbs, shrubs or herbaceous or woody vines that lack stellate trichomes; mostly plurifoliate sympodial unit structure, many compound leaved species | **Regmandra (12)** |  |  | Herbs and shrubs, leaves mostly deeply lobed to compound, thick and somewhat succulent, corollas mostly rotate, stigmas expanded and clavate, stone cells present | Pacific coastal deserts and dry habitats (incl. lomas) of Peru and Chile |
|  |  | **VANans** |  | **Valdiviense (1)** | Woody plant with cup-shaped pedicel bases (insertion), berries red at maturity | Chile |
|  |  |  |  | **African non-spiny (ANS, 14)** | Woody shrubs or vines, mostly climbing, with unbranched or dendritic hairs, plurifoliate sympodial units, terminal inflorescences, and mostly deeply stellate purple corollas | Africa & Madagascar |
|  |  |  |  | **Normania (3)** | Herbs and shrubs with inflated calyces, bilaterally symmetric corollas, heteromorphic anthers with basal modifications (horn-like projections), unbranched internodal inflorescences | Macaronesia, Spain and Mediterranean North Africa |
|  |  |  |  | **Archaeosolanum (8)** | Soft-wooded shrubs with highly variable leaves (simple to deeply lobed) with winged petioles, rotate | Australia and the South Pacific (New Guinea, |
|  |  |  |  |  | purple corollas, loosely erect anthers on relatively long filaments, and berries with many stone cells | Australia, Tasmania, New Zealand) |
|  |  | **DulMo** |  | **Dulcamaroid (43)** | Mostly subwoody to woody (often scandent) shrubs, some climbing with twining petioles, pedicels with distinctive cup-shaped bases (insertions), stone cells absent | Global |
|  |  |  |  | **Morelloid (79)** | Herbs to shrubs, with internodal or leaf-opposing inflorescences, stone cells absent | Global |
|  |  | **Potato** | Mostly herbs, herbaceous vines, or weakly woody vines | **Herpystichum (10)** | Small herbs or herbaceous vines commonly rooting along nodes with simple to 3-5-foliate leaves, flower buds distinctly onion-shaped and fruits mostly flattened | Lowland rain forest and pre-montane forests from southern Mexico to northern Peru |
|  |  |  |  | **Pteroidea (10)** | Understory herbs, herbaceous vines with adventitious roots and unbranched shrubs with unifoliate sympodial units, simple or pinnate leaves, and axillary inflorescences with small, deeply stellate, fruits often pointed and warty | Lowland rain forest and pre-montane forests from Mexico to tropical South America |
|  |  |  |  | ***S. oxycoccoides* (1)** | Somewhat woody vine with simple leaves lacking pseudostipules, and small red fruits with only a few seeds | Peruvian Andes >3,000m |
|  |  |  |  | **Articulatum (2)** | Somewhat woody vines with paired pseudostipules, with large, often branched inflorescences and winged seeds resembling *Basarthrum* clade but lacking bayonet hairs (2-celled hairs with a short apical cell). | Montane forests >2,500m in Costa Rica, Panama, and northern Colombia |
|  |  |  |  | **Basarthrum (16)** | Somewhat woody vines or lax shrubs simple to pinnately compound leaves with frequent interjected leaflets, subtended by a pair of pseudostipules, all species with bayonet hairs (2-celled hairs with a shorter apical cell), with pendant fruits typically green, striped | Dry to moist mid-elevation habitats in Central & South America |
|  |  |  |  | **Anarrichomenum (12)** | Somewhat woody vines readily root along nodes, simple or compound leaves lacking interjected leaflets, subtended by a single (not paired) pseudostipules at most nodes, all species with bayonet hairs (2-celled hairs with a shorter apical cell), fruits orange to red and seeds winged. | Mid-elevation habitats in Central & South America |
|  |  |  |  | **Etuberosum (3)** | Rhizomatous herbs with pinnately compound leaves with frequent interjected leaflets subtended by paired pseudostipules, with many-branched terminal inflorescences, morphologically similar to the Petota clade members that have tubers, pedicels articulated at or near the base (unlike Petota clade). | Argentina and Chile |
|  |  |  |  | **Tomato (17)** | Herbs to woody vines with pinnate leaves with mostly serrate to serrulate margins, some with interjected leaflets, all subtended by paired pseudostipules, all species with strong vegetative odours (glandular trichomes) and yellow corollas, with most species with distinct apical anther modifications (appendages), fruiting pedicels articulating mostly in the distal part | Colombia, Ecuador, Peru and Chile |
|  |  |  |  | **Petota (113)** | Tuber-bearing herbs mostly with compound leaves with interjected leaflets with entire leaflet margins, subtended by paired pseudostipules, inflorescences always branched and terminal, corollas usually rotate, fruiting pedicels articulating mostly in the distal part, berries green to purple | North, Central & South America |
| **Clade II** |  | ***S. anomalostemon* (1)** |  |  | Small herb with glandular trichomes, simple to 3-foliate leaves, distinct cordate shaped anthers with apical modification (beak) | Dry forests in Southern Peru |
|  |  | **Brevantherum** | Non-prickly, shrubs or trees (herbs) with cylindrical anthers, some with stellate hairs | **Trachytrichium (2)** | Shrubs with simple entire leaves and unbranched inflorescences with deeply stellate white corollas, heteromorphic filaments in some species, fruits green | Brazil |
|  |  |  |  | **Inornatum (5)** | Herbs to small shrubs with simple entire leaves, simple trichomes, and unbranched inflorescences with deeply stellate white corollas, fruits green | Brazilian Atlantic forest and Cerrado |
|  |  |  |  | **Gonatotrichum (7)** | Herbs to shrubs, most species with distinctive 2-celled trichomes with a long apical cell that is bent at a 90 degree angle, inflorescences unbranched with mostly rotate white corollas, heteromorphic filaments in some species, fruits with explosively dehiscent | Southern United States to southern South America |
|  |  |  |  | **Brevantherum (80)** | Trees or shrubs with distinct branching pattern where stem | Native to central and South |
|  |  |  |  |  | forks have an inflorescence or a leaf arising from it (dichasial branching) and rank fishy vegetative odour, with stellate trichomes (or lepidote scales), most species with highly branched inflorescences, many with upright and stout peduncle and pedicels, mostly white corollas, fruits various colours remaining green until just before maturity. | America with many species now found globally |
|  |  | **Geminata** | Non-prickly shrubs or trees with simple entire leaves and rank vegetative odour (burning smell), anthers cylindrical. | **Reductum (2)** | Small shrubs with dendritic hairs, broadly stellate white corollas, and one species with basal anther modifications (sack). | Argentina |
|  |  |  |  | **Geminata (150)** | Shrubs or trees, many with geminate (paired/twinned) leaves, inflorescences mostly leaf-opposed and with deeply stellate white flowers with relatively stout, oblong anthers, and fruits that remain green at maturity | Moist lowland or montane forests from Central to South America |
|  |  | **Cyphomandra** | Non-prickly shrubs, small trees or woody vines with rank vegetative odour (burning smell), plurifoliate or trifoliate sympodial growth, tapered anthers, and with long and mostly branching inflorescences | ***S. graveolens* (1)** | Woody vine with pinnately compound leaves, white broadly stellate corollas, anthers with apical modification (beaks) | Brazil |
|  |  |  |  | **Cyphomandropsis (11)** | Shrubs or small trees with simple entire leaves, many with dendritic trichomes, mostly white corollas, some with stone cells | Mostly dry habitats across South America |
|  |  |  |  | **Pachyphylla (42)** | Shrubs (often lax & spreading) or small trees with distinct branching pattern where stem forks have an inflorescence or a leaf arising from it (dichasial branching), some species with lobed or compound leaves, corollas mostly deeply stellate, anthers with enlarged connectives, some with stone cells | Lowland and pre-montane moist forests from Mexico to South America |
|  |  | **Wendlandii-Allophyllum** | Prickly to non-prickly shrubs, herbs and woody vines with tapered anthers | **Allophyllum (4)** | Shrubs or herbs lacking prickles, with distinct branching pattern where stem forks have an inflorescence or a leaf arising from it (dichasial branching), leaves covered in idioblasts containing crystal sand (sand-punctate), inflorescences simple, few-flowered and relatively short, corollas broadly stellate or rotate, white or greenish white, some species with stone cells | Lowland moist and montane forests of Central and South America |
|  |  |  |  | **Wendlandii (9)** | Woody vines with mostly broad-based recurved prickles and with large and generally many-branched inflorescences, many species with lobed or compound leaves, heteromorphic filaments, or swollen calyces in fruit. | Mexico, and Central and South America |
|  |  | **Nemorensis** | Prickly vines, shrubs or herbs, andromonoecious, with tapered anthers | **Nemorensis (4)** | Woody or herbaceous vines or herbs with mostly broad-based recurved prickles, sympodial units mostly difoliate geminate, leaves usually lobed or compound, inflorescences simple and leaf-opposed with flowers with deeply stellate corollas. | South America |
|  |  | **Leptostemonum** | Prickly shrubs, trees, woody vines or herbs with stellate trichomes, | ***S. polygamum* (1)** | Dioecous shrub or small tree with some needle-like prickles, with simple entire leaves, deeply stellate 5-6-merous white corollas, white, cylindrical anthers, female | Caribbean |
|  |  |  | difoliate (geminate or non-geminate) sympodial units dominate, leaves simple or lobed, mostly andromonoecious, simple inflorescences dominate, anthers tapered |  | flowers lacking developed anthers and with large, forked stigmas, and pubescent orange fruits subtended by large leafy calyx lobes. |  |
|  |  |  |  | **Lasiocarpa (12)** | Shrubs or small trees with needle-like prickles, with mostly repand leaves and simple inflorescences, mostly broadly stellate corollas and orange fruits covered in stellate usually glandular trichomes. | Global (mostly Northern and Western South America, with two species in Asia and the Pacific) |
|  |  |  |  | **Acanthophora (22)** | Shrubs and herbs with deeply stellate corollas characterised by having a mix of stellate and simple trichomes on lower leaf surfaces but only simple trichomes on stems and upper leaf surfaces. | Disturbed and open habitats in South America (mostly eastern Brazil) |
|  |  |  |  | **Gardneri (10)** | Slender stemmed shrubs and herbs mostly with needle-like prickles, small simple (mostly unlobed) leaves, short and laterally directed inflorescences, deeply stellate corollas, and berries with somewhat accrescent calyces covering less than half of the fruit. | Dry habitats of eastern to central Brazil, Caribbean, Mexico and Central America, and northern Peru |
|  |  |  |  | **Thomasiifolium (9)** | Shrubs or woody vines mostly with broad-based recurved prickles, some species with glandular-stellate trichomes, stems often with short internodes and leaves grouped at the apex, inflorescences unbranched, corollas purple, some bilaterally symmetric, fruits either large and densely pubescent with large seeds, or small and glabrous with accrescent calyces that cover less than half of the fruit. | Eastern Brazil |
|  |  |  |  | **Erythrotrichum (35)** | Shrubs, woody vines or small trees covered in broad-based recurved prickles, with stellate glandular trichomes, trichomes dense and often reddish-brown in colour, fruits covered in stellate trichomes. | Tropical South America |
|  |  |  |  | **Sisymbriifolium (2)** | Shrubs or herbs with dense needle-like prickles, leaves deeply lobed (nearly pinnate), corollas broadly stellate to nearly rotate, white, fruits red with appressed spiny calyces that spread open at full maturity. | Dry habitats of South America |
|  |  |  |  | **Crinitum (23)** | Trees, large shrubs or woody vines with scattered broad-based prickles and large flowers with bilaterally symmetric purple corollas and long, heteromorphic anthers, berries large and hardened, oxidize black when cut open, with distinctly swollen calyces. | Mexico and South America |
|  |  |  |  | **Androceras (16)** | Herbs densely covered in needle-like prickles, with bilaterally symmetric corollas with heteromorphic anthers, dry fruits surrounded by accrescent spiny calyces. | Southern USA and Mexico |
|  |  |  |  | ***S. campechiense* (1)** | Small shrub covered in needle-like prickles, corollas rotate, fruits with appressed calyces lacking pedicel articulation. | Dry forests of Mexico, and Central and South America |
|  |  |  |  | **Carolinense (14)** | Rhizomatous herbs and shrubs covered in needle-like prickles, some species with tubers, Corollas rotate, mostly purple, fruits green to yellow mottled, some with appressed calyces. | Open habitats of Southeastern USA and Bolivia, Argentina and Paraguay |
|  |  |  |  | **Bahamense (3)** | Shrubs and trees with some needle-like prickles, deeply stellate corollas, anthers with stellate trichomes on the adaxial surface, and small juicy red or black fruits on strongly recurved fruiting pedicels. | Caribbean |
|  |  |  |  | **Micracantha (14)** | Scandent shrubs and woody vines that climb using broad-based recurved prickles with unbranched inflorescences, deeply stellate (mostly white) corollas, and mostly orange or red fruits. | Florida to Bolivia, including the Caribbean |
|  |  |  |  | **Asterophorum (4)** | Tuber-bearing shrubs with difoliate geminate sympodial growth, with broad-based prickles, simple leaf-opposing inflorescences with broadly stellate white corollas, and fruit with appressed calyces covering at least half of the fruit. | Brazilian Atlantic forest |
|  |  |  |  | ***S. multispinum* (1)** | Rhizomatous small shrub covered in needle-like prickles, with white broadly stellate corollas, and yellow berries with prickly accrescent calyces that cover less than half of the fruit. | Dry habitats of Argentina, Brazil and Paraguay |
|  |  |  |  | **Torva (55)** | Shrubs, small trees or woody vines with mostly branched (and stout) inflorescences, with some needle-like or occasionally broad-based | Global (mostly tropical Americas, with a |
|  |  |  |  |  | prickles, usually broadly stellate corollas, and fruits on stout pedicels often held upright with, most species with mucilaginous pulp. | few members in Asia) |
|  |  |  |  | *S. euacanthum* (1) | Herb covered in needle-like prickles and stellate glandular trichomes, with broadly stellate white corollas, and dehiscent fruits lacking mesocarp and ‘exploding’ when ripe (probably from tension in the exocarp) with appressed calyces that cover more than half of the fruit. | Dry habitats of subtropical Argentina. |
|  |  |  |  | *Elaeagnifolium* (5) | Rhizomatous shrubs or herbs with various degrees of both broad-based recurved and needle-like prickles, weakly bilaterally symmetric purple corollas with heteromorphic anthers and dry fruits, some with lepidote scales. | Deserts and dry habitats in North and South America |
|  |  |  |  | **Eastern Hemisphere Spiny (EHS, 336)** | Morphologically diverse group of shrubs, herbs, woody vines or small trees), mostly densely prickly (broad-based recurved and/or needle-like), some rhizomatous, many with appressed calyces that cover more than half of the fruit. Most prickly Solanum species outside the Americas are members of this clade | Africa, Madagascar, Asia, Australia, Pacific |

Ten further traits show evolutionary lability with 50-100 transitions across the genus, including specialised underground organs, prickles, trichome structure, leaf type, inflorescence position and branching, sexual system, stamen heteromorphism, presence of trichomes on mature fruit, and fruiting calyx modifications (Table 1; Figs. 5-6). Underground storage organs have evolved at least four times independently, while rhizomes have been gained 30 times and lost at least 27 times across *Solanum* (Fig. 5A; SI Table 3). Prickles have a single origin in *Solanum*, with needle-like acicular prickles modelled as ancestral followed by several switches between needle-like and broad-based prickles, including multiple losses of prickles (Fig. 5B). Prickles are found exclusively in clades with stellate trichomes, except for the Wendlandii and Nemorense clades that lack stellate trichomes but have prickles (Fig. 5B-C). Stellate trichomes have evolved twice in *Solanum* from dendritic trichomes and have been lost several times (Fig. 5C; SI Table 3). Most frequent shifts include changes between dendritic to simple trichomes (Fig. 5C; SI Table 3). Leaves in *Solanum* are modelled to have been compound ancestrally with frequent switches to simple and then to lobed (Fig. 5D).

**Fig. 5.**
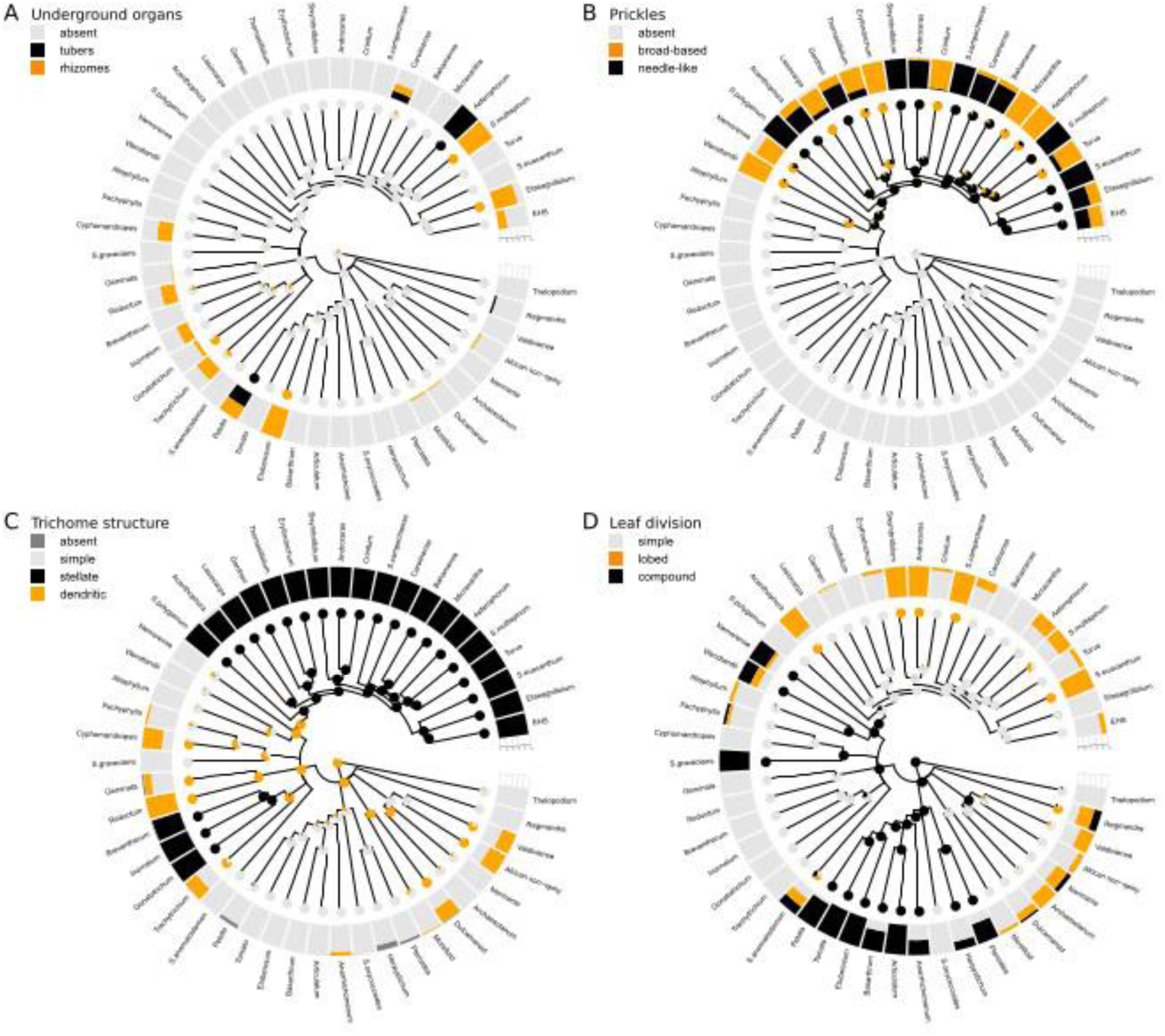
Evolution of labile morphological traits (vegetative) in *Solanum* with 50-100 transitions based species-level analysis using stochastic character mapping. **A**, Specialised underground organs; **B**, Prickles; **C**, Trichome structure; and **D**, Leaf division. Results from the best model are shown for each character (see Table 2 for details) based on 200 simulations. The topology used for mapping was derived from a supermatrix phylogeny with nine loci (two nuclear and seven plastid loci; Gagnon & al., 2022) with 725 species sampled and coded for each trait (60% of all species). All major and minor clades are labelled; tips reflect the crown nodes of each minor clade. Piecharts indicate likelihood of modelled ancestral states along the nodes, and frequency bars (tips) reflect proportion of species sampled within each clade with each state.

**Fig. 6.**
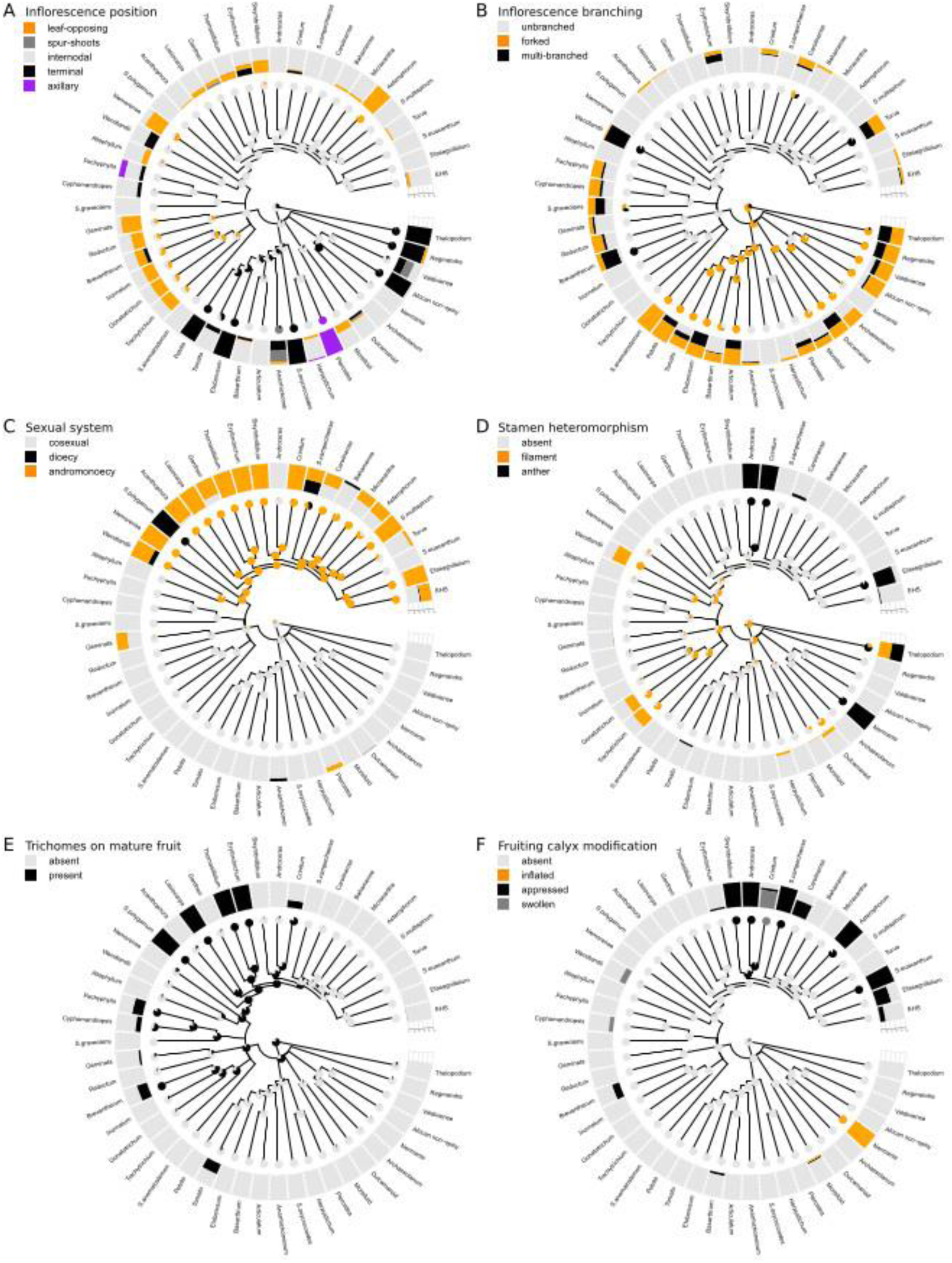
Evolution of labile morphological traits (reproductive) in *Solanum* with 50-100 transitions based species-level analysis using stochastic character mapping. **A**, Inflorescence position; **B**, Inflorescence branching; **C**, Sexual system; **D**, Stamen heteromorphism; **E**, Trichomes on mature fruits; and **F**, Fruiting calyx modifications. Results from the best model are shown for each character (see Table 2 for details) based on 200 simulations. The topology used for mapping was derived from a supermatrix phylogeny with nine loci (two nuclear and seven plastid loci; Gagnon & al., 2022) with 725 species sampled and coded for each trait (60% of all species). All major and minor clades are labelled; tips reflect the crown nodes of each minor clade. Piecharts indicate likelihood of modelled ancestral states along the nodes, and frequency bars (tips) reflect proportion of species sampled within each clade with each state.

Internodal, forked inflorescences are modelled as ancestral in *Solanum* with repeated shifts in both inflorescence position and branching across the phylogeny where multi-branching inflorescences have been frequently gained and lost (Fig. 6A-B). Terminal branched inflorescences are more common in Grade I while unbranched internodal inflorescences dominate in the large Leptostemonum clade (Fig. 6A-B). The ancestral sexual system in *Solanum* is cosexuality (i.e., hermaphroditism) with a minimum of 17 independent origins of andromonoecy and >60 reversals back to cosexuality across *Solanum* (Fig. 6C). Dioecy has evolved at least six times in *Solanum*, once from cosexual ancestors and five times from andromonoecy (Fig. 6C; SI Table 3). Heteromorphic anthers have evolved at least 13 times in *Solanum* and have rarely been lost (Fig. 6D). Heteromorphic filaments are modelled ancestral in *Solanum* with frequent losses and re-gains; heteromorphism is generally present in either filaments or anthers, except for the Thelopodium clade where all species have both heteromorphic filaments and anthers (Fig. 6D). Trichomes on mature fruits are modelled ancestral in *Solanum* with frequent losses and re-gains (Fig. 6E). Appressed fruiting calyces are common in *Solanum* with 48 independent origins, most of which are found in the Leptostemonum clade (Fig. 6F; SI Table 3). Inflated calyces have evolved minimum of five times (Fig. 6F; SI Table 3).

Five floral and fruit traits show 10-42 changes across *Solanum* indicating relatively conserved evolution, including corolla bilateral symmetry, anther shape, pedicel articulation, fruit type and presence of stone cells in fruit (Table 1; Fig. 7). Bilaterally symmetric corollas have evolved seven times independently with frequent losses (Fig. 7A; SI Table 3). Tapered anther shape has evolved 6 times independently from an ancestral cylindrical shape, and cordate anthers have two independent origins (Fig. 7B; SI Table 3). Fruiting pedicels dehisce basally in most *Solanum* but articulation towards the distal end has evolved in the Petota-Tomato clade and has been lost twice (Fig. 7C; SI Table 3). Dry dehiscent berries have evolved at least 13 times in *Solanum* and have been rarely lost once gained (Fig. 7D; SI Table 3). Stone cells in fruits are modelled as ancestral in *Solanum* with frequent losses (Fig. 7E).

**Fig. 7.**
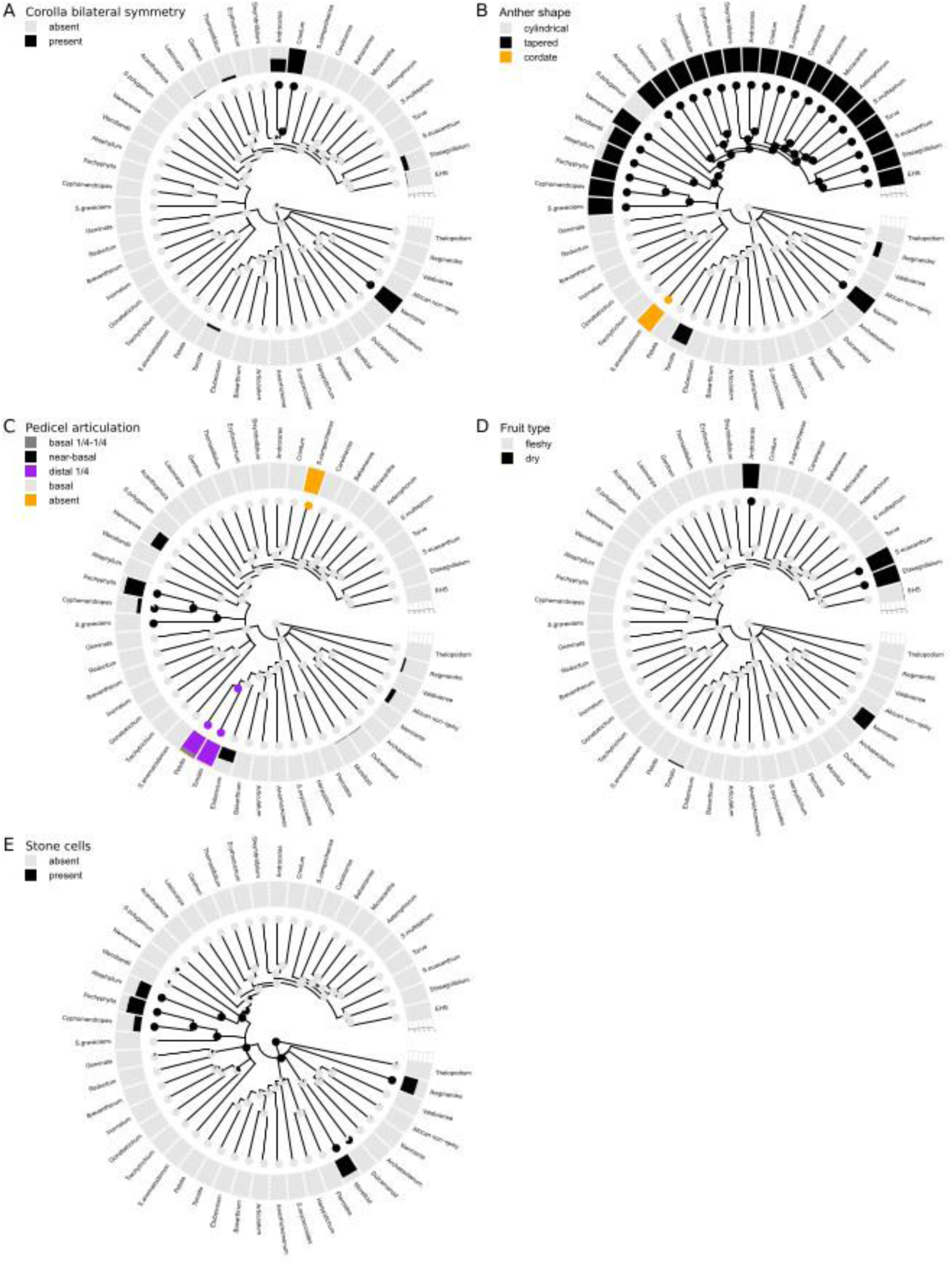
Evolution of conserved morphological traits in *Solanum* with <50 transitions based species-level analysis using stochastic character mapping. **A**, Corolla bilateral symmetry; **B**, Anther shape; **C**, Pedicel articulation; **D**, Fruit type; and **E**, Stone cells. Results from the best model are shown for each character (see Table 2 for details) based on 200 simulations. The topology used for mapping was derived from a supermatrix phylogeny with nine loci (two nuclear and seven plastid loci; Gagnon & al., 2022) with 725 species sampled and coded for each trait (60% of all species). All major and minor clades are labelled; tips reflect the crown nodes of each minor clade. Piecharts indicate likelihood of modelled ancestral states along the nodes, and frequency bars (tips) reflect proportion of species sampled within each clade with each state.

Four highly conserved traits show <10 changes across *Solanum*, including presence of pseudostipules, enlarged anther connectives, anther modifications, and pedicel insertion type (Table 1; Fig. 8). Pseudostipules have evolved multiple times but only in two clades (Fig. 8A). Enlarged anther connectives have evolved only once but have been lost frequently (Fig. 8B). Other anther modifications have evolved five times independently in *Solanum*, all outside the large Leptostemonum clade (Fig. 8C; SI Table 3). Cup-shaped pedicel insertion has evolved twice (Fig. 8D).

**Fig. 8.**
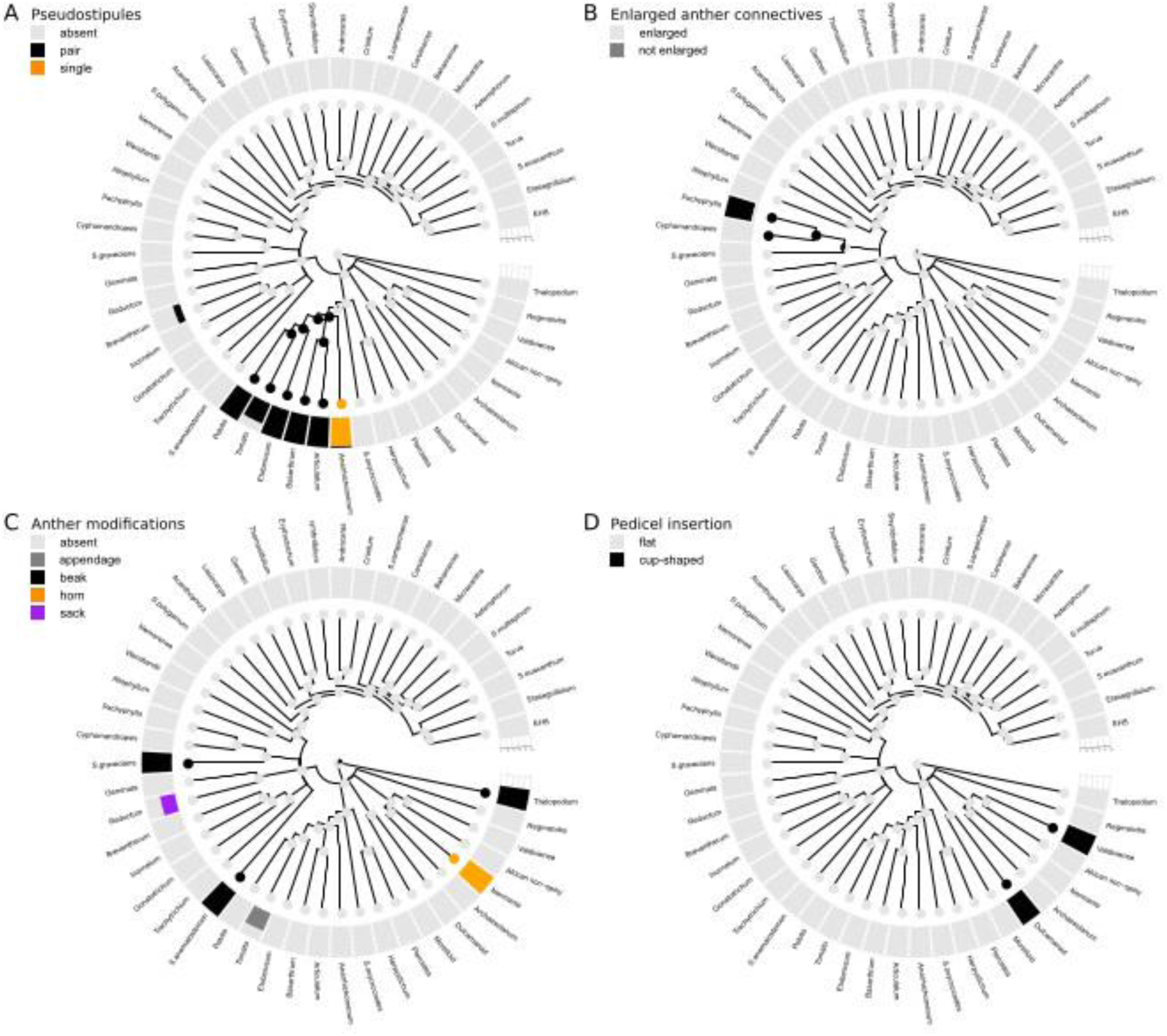
Evolution of the most conserved morphological traits in *Solanum* with <10 transitions based species-level analysis using stochastic character mapping. **A**, Pseudostipules; **B**, Enlarged anther connectives; **C**, Anther modifications; and **D**, Pedicel insertion. Results from the best model are shown for each character (see Table 2 for details) based on 200 simulations. The topology used for mapping was derived from a supermatrix phylogeny with nine loci (two nuclear and seven plastid loci; Gagnon & al., 2022) with 725 species sampled and coded for each trait (60% of all species). All major and minor clades are labelled; tips reflect the crown nodes of each minor clade. Piecharts indicate likelihood of modelled ancestral states along the nodes, and frequency bars (tips) reflect proportion of species sampled within each clade with each state.

## Discussion

### Morphological definition of clades

Identifying morphological traits that characterise groups has been one of the principal aims in taxonomy since its origin (Humphreys & Linder, 2009). Views on the importance of vegetative versus reproductive traits in defining genera has changed over the centuries (e.g., Stevens, 1997). Linnaeus (1735, 1753) used a so-called ‘sexual system’ based solely on numbers of male and female parts in flowers for classifying angiosperms into smaller units. Both Linnaeus’ contemporaries (e.g., Adanson, 1764) and subsequent taxonomists attempted more ‘natural’ classifications, based on a wider selection of characters derived from all plant parts. Pheneticists continued this tradition by advocating the use of all characters with equal weight, in contrast to cladists who focused on the use of shared derived traits in defining groups (Williams & Ebach, 2020). A focus on a few obvious (often reproductive) traits has led to the recognition of many morphologically distinct small genera (Humphreys & Linder, 2009; Stevens, 1997). Molecular phylogenetic analyses have, however, revealed many genera (or higher rank taxa) to be para-or polyphyletic, due in large part to the convergent evolution (i.e., evolutionary lability) of many floral and fruit traits used to define these groups prior to the advent of DNA sequence data (e.g., Orejuela & al., 2017; Bogarín & al., 2019; Koenen & al., 2020; Appelhans & al., 2021).

Traits that define clades within big genera have remained elusive in part because of the logistical difficulty of studying large groups (Minelli, 2016). *Solanum* is an example of such an unwieldy group, where the efforts of a large and collaborative team of taxonomists, field botanists, and molecular systematists now enable us to provide a broad overview of morphological diversity across the genus at species-level for the first time. Our analysis is a major step forward in quantifying morphological diversity in *Solanum*, one of the largest plant genera across the tree of life (Minelli, 2016). The analyses presented here allow us to morphologically define the informally named clades of *Solanum*, as presented in Table 2. The most conserved traits are clearly the most useful in defining clades, but even the most labile traits prove useful in defining particular clades when rare states are expressed and if used in combination (e.g., axillary inflorescences in Pteroidea clade with unifoliar sympodial unit structure, or presence of both yellow corollas and pseudostipules in Tomato clade). Such fine-tune understanding of variation (or lack of) enables us to confidently place all currently accepted species of *Solanum* into the informal clades with only a few exceptions (SI Table 4). Lastly, the morphological definition of clades is also a powerful identification tool, and that in mind, the underlying data has been used to construct an online interactive multi-access format key to the groups (Solanum Key Consortium, 2022).

### Plant structure, roots and leaves

Plant structure related traits have some of the highest transition rates in *Solanum* with repeated shifts between sympodial growth patterns and herbaceous and woody growth forms (Figs. 1A-F, 4A-B). All herbaceous vines and epiphytes in *Solanum* are found in Grade I except for a single truly herbaceous vine species found in the Nemorense clade (*S. hoehnei*, Fig. 4A), while woody growth forms predominate in Clade II. Shifts between growth forms are most prevalent in Grade I where all seven states are observed (Fig. 4A). Growth pattern in *Solanum* is sympodial, where stems are composed of a series of concaulescent branches where flowering marks the end of the main shoot development and axillary buds continue plant growth (Danert, 1958, 1970; Child, 1979). Thus, changes in the sympodial unit structure lead to variation in plant structure in general as well as inflorescence position (Danert, 1958, 1970; Child, 1979). Genetic control of the number of leaves per sympodial unit is known to involve the tomato self-pruning (sp) locus: the sympodial units of sp mutants are terminated early and have fewer leaves (Paran & van der Knaap 2007). The great lability observed here for both traits linked to plant structure suggest that growth form has played a key role in diversification in *Solanum.* Sympodial growth structure is perhaps the closest trait in the genus to the concept of ‘key innovation’ seen in other large plant genera, such as insular woodiness in *Begonia* L. (Kidner & al., 2016) or succulence and the linked evolution of CAM photosynthesis in *Euphorbia* L. (Anest & al., 2021; Horn & al., 2012, 2014). Growth form variation in *Solanum* is often underappreciated, perhaps due to the difficulty of capturing it in photos or in herbarium specimens.

In comparison to the high evolutionary lability of aboveground growth form, the evolution of specialised underground organs is less, albeit still evolutionarily labile. Our analyses show repeated gains and losses of rhizomes and four origins of underground storage organs in *Solanum*; three of the transitions have resulted in stem tubers (Petota, Carolinense, and Asterophorum clades, the latter two members of Leptostemonum) while the fourth has given rise to a swollen caudex (*S. montanum* Cav., Regmandra clade; Fig. 5A). Although stem tubers are often associated with herbaceous species (e.g., potatoes), in the Leptostemonum clade they are found in woody shrubs (members of the Asterophorum clade; Fig. 5A). Rhizomes appear evolutionarily more labile compared to tubers, and the relationship between rhizomes and tubers bears further investigation as both are largely found in the same clades (Fig. 5A; Table 2).

### Plant defence: Glands, trichomes and prickles

Glandular trichomes are involved in herbivore resistance and in *Solanum* contain a variety of different chemical compounds (Weinhold & Baldwin, 2011; Glas & al., 2012; Fan & al., 2019). The high evolutionary lability of glandular trichomes observed in *Solanum* with frequent gains and losses is not surprising, considering many species show infraspecific variation in glandular pubescence between populations and individuals (e.g., *S. nigrum* L.*, S. retroflexum* Dunal, *S. villosum* Mill.; Manoko, 2007; Särkinen & al., 2018). Glandular trichomes are most prevalent in simple-haired clades; only seven (minor) clades have stellate glandular trichomes (Fig. 4C). Morphological variation in simple glandular trichomes has been studied extensively in the Tomato clade where seven distinct types have been identified (types I-VII; Simmons & Gurr 2004); each type has distinct genetic control (Schilmiller & al., 2009, 2012; Zhang & al., 2015; Chang & al., 2018; Chalvin & al., 2020) and potentially distinct functions. Some of these with single-celled glandular tips contain high amounts of acyl sugars and are involved in insect resistance (types I and IV; e.g., Weinhold & Baldwin, 2011), while trichomes with multicellular glandular tips secrete terpenes with various functions (type VI; e.g., Glas & al., 2012; Fan & al., 2019). The widespread occurrence of glandular trichomes across the clades of *Solanum* provides an ideal system with which to test homology of genetic and developmental pathways beyond the Tomato clade.

Prickles in *Solanum* are epidermal in origin and are thought to be modified multicellular stellate trichomes with layers of elongate and lignified cells (Seithe, 1962, 1979; Whalen 1984). Our results show that the single origin of prickles corresponds to the evolution of stellate trichomes in the Leptostemonum clade (Fig. 5B-C), supporting the view that prickles have originated from stellate trichome type. The common origin of stellate trichomes and prickles can be observed in some Leptostemonum species (e.g., *S. barbisetum* Nees, *S. myoxotrichum* Baker and *S. schumannianum* Dammer, all within the EHS clade) where stellate trichomes on young stems develop lignified stalks and become prickle-like with an apical stellate trichome (Vorontsova & Knapp 2016; Aubriot & Knapp, 2022). Previous studies have proposed glandular trichomes to be involved in prickle development (Pandey & al., 2018) but these hypotheses remain unsupported based on our findings and other studies (Zhang & al., 2021). Once gained, prickles have gained diverse morphologies within *Solanum* and have been lost several times, where many of the losses are associated with domestication (e.g., *S. sessiliflorum*, *S. stramonifolium*, *S. quitoense*, *S. aethiopicum*, *S. macrocarpon*, *S. melongena*; Lester & Thitai 1989; Whalen & al., 1981). Variation in prickle type and density observed within species and individuals shows that prickle expression is highly labile; prickles and trichomes are observed to be denser/exclusive to juvenile individuals in some species (e.g., Vorontsova & Knapp, 2016) while in others prickly and non-prickly stems can occur on the same plant (e.g., *S. elaeagnifolium* Cav.; Knapp & al., 2017). Prickles have been shown to be under simple dominant inheritance genetically (Lester & Thitai, 1989), and QTLs and candidate genes responsible for prickle formation have been identified (Portis & al., 2015; Pandey & al., 2018; Miyatake & al., 2020; Qian & al., 2021; Zhang & al., 2021). Studies exploring the development of prickles through the life of a plant are needed, and the ecological function of prickles could be explored via analyses of climatic correlates.

Trichome structure, in contrast, is less labile and defines some of the major clades in *Solanum* (Fig. 5C; Dunal, 1852; Seithe, 1962, 1979; Roe, 1971, 1972). Stellate trichomes have arisen twice in Clade II (Fig. 5C). Dendritic trichomes, which never occur in clades with stellate trichomes, are modelled as ancestral in *Solanum* and have been lost several times (Fig. 5C). Seithe (1962, 1979) observed simple trichomes on seedling leaves of species that subsequently developed dendritic or stellate trichomes, leading her to suggest that both were derived from simple trichomes. Our results, which model dendritic trichomes as ancestral, do not support this view. New studies, focusing on the phylogenetic context of trichome development, will be needed to shed light on our seemingly counterintuitive result. There is more morphological variation in trichome structure than displayed by the broad categories used in our analyses, with several losses and modifications of stellate trichomes within clades. For instance, the 2-celled geniculate trichomes in the Gonatotrichum clade with a short basal and longer apical cell are thought to represent reduced stellate trichomes (Seithe, 1979; Stern & al., 2013). This is an area of great promise for future research in developmental genetics.

### Leaves

Compound (i.e., deeply pinnatifid) leaves are modelled as ancestral in *Solanum,* with frequent shifts to simple (entire) leaves (Fig. 5D). The trait model predicts, surprisingly, direct shifts from compound to simple leaves without an ‘intermediate’ step of leaf lobbing (SI Table 3), as can be seen in particular clades with both simple and compound leaved species, but lacking species with ‘intermediate’ lobed leaves (e.g., Herpystichum, Pteroidea, Anarrhichomenum, Basarthrum, and Petota; Fig. 5D). In these clades, simple leaves may represent reduced compound leaves where only a singe leaflet remains, as this can be observed in some tomato mutants (Berger & al., 2009). Transitions to lobed leaves are modelled to be via simple entire leaves (SI Table 3), and such transitions can be observed in Clade II, where some species show simple, lobed, and compound leaves in a single shoot (e.g., Wendlandii and Pachyphylla clades: *S. wendlandii* Hook.f. or *S. pendulum* Ruiz & Pav., respectively). All species in Leptostemonum have either simple or lobed leaves (some deeply lobed, e.g., *S. sisymbriifolium* Lam., coded as lobed), and change in shape from lobed to entire has been documented in the Torva clade with developmental maturity of the shoot (Roe, 1966). Our results disagree with previous studies on leaf division patterns across *Solanum* where leaves were modelled as ancestrally simple and entire (Geeta & al., 2012), the difference in our results likely due to increased species sampling and the fact that we collapased many nodes along the backbone of *Solanum* due to the high discordance and topological uncertainty based on phylogenomic sampling (Gagnon & al., 2022). Genetic studies show upregulation of KNOX genes in both compound and lobed leaves in *Solanum* but not in simple leaves (Hagemann & Gleissberg, 1996; Bharathan & al., 2002; Efroni & al., 2010). Exploring the evolution of genes linked to leaf patterning would be interesting along major clades of *Solanum*, and including in lineages where considerable polymorphism within individual plants can been observed, following approaches in Brassicaceae (Vlad & al., 2014; Streubel & al., 2018; Nikolov & al., 2019).

### Floral traits

Two traits linked to pollinator attraction, corolla shape and colour (Møller, 1995; Gómez & al., 2008, 2016; Muchhala & al., 2014; Reverte & al., 2016; Moré & al., 2020), are shown to be some of the most evolutionarily labile traits in *Solanum* (Fig. 4D-E). Variation in corolla shape in *Solanum* is driven by the amount of interpetalar tissue present between lobes, and changes seem frequent throughout the genus highlighting the need to explore pollinator-linked trait variation in buzz-pollinated plant groups. Future studies should aim to quantify corolla shape to include corolla lobe orientation to enable more detailed studies of trait evolution at species-level.

Shifts between purple and white dominate in *Solanum* with >66 gains of purple corollas (Fig. 4E). Transitions to yellow are rare with a minimum five independent origins of yellow corollas, twice from white ancestors similar to results from Antirrhineae (Orobanchaceae) where transitions from purple to yellow have been found to have occurred via white (i.e., ‘loss’ of previous pathway; e.g., Ellis & Field, 2006). Three transitions in *Solanum*, however, indicate direct shifts from purple to yellow (Fig. 4E); these nodes merit further investigation. The strength and brightness of yellow is not homogenous across *Solanum*, and it is possible that this colour arises from different pathways and is not homologous. Genetic control of corolla colouration (e.g., Gates & al., 2018) has not been studied in *Solanum*, but flavonoids (anthocyanins: purple; flavones: yellow) and carotenoids (yellow) are components of corolla colouration in other genera of Solanaceae (*Petunia* Juss.: Berardi & al., 2021; *Iochroma* Benth.: Berardi & al., 2016, Larter & al., 2019; *Nicotiana* L.: McCarthy & al., 2017). We did not record the presence of multiple colours in some species (e.g., corollas with shiny central ‘eyes’ at the base of the corolla with green, yellow, purple, or black colouration) but this merits further investigation. Studies on *Antirrhinum* L. have shown that even single mutations altering corolla colour and reflectance can affect bee behaviour (Comba & al., 2000; Dyer & al., 2006, 2007).

Our results indicate that the general floral ‘bauplan’ in *Solanum* has changed several times independently despite the relative morphological homogeneity in *Solanum* flowers based on the buzz-pollination syndrome with poricidally dehiscent anthers arranged in a central cone. Changes include a minimum of 13 origins of heteromorphic anthers, eight origins of heteromorphic filaments (Fig. 6D), seven origins of bilaterally symmetric corollas (Fig. 7A), and five independent origins of anther modifications (Fig. 8C). Bilaterally symmetric corollas have evolved largely in clades with heteromorphic anthers (Leptostemonum; Figs. 4C & 7B; Lester & al., 1999; Bohs & al., 2007; Knapp, 2002b; Knapp, 2010). The link between corolla and stamen zygomorphy has been well-documented and has evolved multiple times in Solanaceae (Robyns, 1931), where heteromorphic anthers appear to be precursors to bilaterally symmetric corollas (Zhang & al., 2017). Genes controlling corolla bilateral symmetry (and heteromorphic anthers) in *Solanum* have not yet been identified, but the trait is likely to affect pollination by increasing pollinator specificity and efficiency (Jesson & Barrett, 2002, 2005; Fenster & al., 2009). Enlarged anther connectives have evolved once in *Solanum* (Fig. 7C) again affecting pollination as they are involved in the perfume bee pollination in the Pachyphylla clade (Sazima & al., 1993; Bohs, 1994; Cocucci, 1996, 1999; Falcão & Stehmann, 2018). Taken together, our results indicate continuous shifts in pollination related floral traits in a buzz-pollinated genus, and the drivers of this variation should be explored further in species-level studies.

### Fruit morphology

*Solanum* is known for variation in fruit shape, size, colour, and texture (Fig. 1Y-AD; Knapp, 2002c), and fruit traits have been suggested to have been important in the diversification of the EHS clade (Echeverría-Londoño & al., 2020) likely due to their function in dispersal adaptations and consequent ability to colonize new areas and habitats (e.g., Cipollini & al., 2002; Martine & al., 2019). Colour is the most evolutionarily labile fruit character in *Solanum*. Fleshy *Solanum* berries are known to be dispersed by a wide variety of vertebrates, including birds, bats, and small rodents (Symon, 1979a; Cipollini & Levey, 1997a, 1997b, 1997c; Knapp, 2002a), and variation in fruit colour could be linked to changes in dispersal strategies. Green fruits are the most common in *Solanum* (275 species sampled in the phylogeny; SI Table 2), but the commonness of green fruits may be an artefact of late maturing fruits in some groups, where fully mature fruits are rarely collected. A great example of such fruits are the green fruits of the Geminata and Brevantherum clades that go through a rapid change in colour from green to yellow, orange, red or purple when they are mature and are quickly consumed by frugivores such as bats (Knapp, 2002a; Tovar & al., 2021). Yellow (157 spp.) and red fruits (122 spp.) are the second most common fruits in *Solanum*, followed by purple (77 spp.), orange (65 spp.) and white (29 spp.; SI Table 2). Flavonoids, particularly anthocyanins, are responsible for purple, blue, and red fruit colours in Solanaceae, whereas carotenoids are typically responsible for yellow, orange, and red fruits (Gonzali & Perata, 2021). Because we only scored the external peel colour and did not code flesh colouration, future studies should expand on the distinct colour pathways potentially present in fruit flesh and peel in some species such as *S. lycopersicum* (Tomato clade: Ronen & al., 1999; Bovy & al., 2007; Gonzali & Perata, 2021), *S. betaceum* (Pachyphylla clade; Acosta-Quezada & al., 2015), and *S. melongena* (EHS clade; Jiang & al., 2016).

Presence of trichomes on mature fruits and fruiting calyx modifications (Table 2; Fig. 6E-F) are evolutionarily labile in *Solanum*; our results agree with Deanna & al. (2019) showing a stepwice and directional evolution from non-accrescent to accrescent (appressed) and inflated but indicate a second path from non-accrescent directly to inflated (SI Table 3). Fruiting calyx modifications (Table 2; Fig. 6E-F) enable physical dispersal mechanisms, for example where prickly accrescent fruiting calices around dry berries act as trample-burrs or censer fruits where seeds shake out of the open end of the accrescent calyx with plant movement (e.g., *S. euacanthum*, Elaeagnifolium, Androceras and EHS clades; Symon, 1979a; Martine & al. 2019).

The remaining fruit traits in *Solanum* are phylogenetically conserved (Fig. 8A-F; pedicel insertion, pedicel articulation, fruit type, and presence of stone cells) and are useful for defining groups (Table 2). Stone cells (i.e., sclerotic granules or brachysclereids), hard inclusions found in the fleshy portion of some *Solanum* berries, have evolved a minimum of four times (Fig. 8F). Stone cells are derived from sclerenchyma with massively enlarged cell walls and vary in their number, size, shape, and colour (Bitter, 1911, 1914; Symon, 1994; Knapp, 2002c; Särkinen & al., 2018; Knapp & al., 2019). Their function remains unclear, but interestingly, in the Morelloid clade, where stone cells are common, species cultivated for their berries lack them completely (e.g., *S. scabrum* Mill., see Särkinen & al., 2018).

There is complex variation in fruit texture, shape, and size across *Solanum* not captured by our analyses that likely contributes to dispersal strategies (e.g., Cipollini & al., 2002). In our study, the single category of fleshy berries included small, soft, and juicy (e.g., tomatoes), large and spongy (e.g., eggplant), large and woody (e.g., *S. sycophanta* Dunal), and explosively dehiscent fleshy berries (e.g., all species of the Gonatotrichum clade; Knapp, 2001, 2002c; Stern & al., 2013; *S. mellobarretoi* Agra & Stehmann in the Leptostemonum clade; Agra & Stehmann, 2016). Phenotypic variation of fruit texture, shape, and size will be challenging to quantify in all species but will be needed to enhance understanding of diversity across *Solanum*. Methods developed for tomatoes (e.g., van der Knaap & al., 2014) applied across the genus may reveal new characters and character combinations for developmental study. Links between the evolution of fruit type, size, colour, and texture combined with fruiting calyx modifications warrant further exploration because they all affect fruit and seed dispersal.

### Sexual system

Cosexuality, with all flowers having both male and female parts (Cardoso & al., 2018), is ancestral in *Solanum* (Fig. 6C). Dioecy is relatively rare in *Solanum* being found in 15 species sampled so far in the molecular phylogeny, but has evolved at least six times independently, directly from a cosexual (i.e., hermaphroditic) ancestor in one case (Anarrhichomenum clade; Fig. 6C) and from andromonoecious ancestors in others (e.g., Martine & al., 2006, 2009). Plants with dioecious or plastic sexual systems continue to be discovered in *Solanum* (e.g., Knapp & al., 1998; Martine & al. 2016; McDonnell & al., 2019) with a total of 22 dioecious species known in *Solanum* currently (Fig. 3D). Andromonoecy as coded here is common in *Solanum* and is modelled as ancestral in the Leptostemonum clade and its sister lineage with multiple gains and losses (Fig. 6C). There is large variation within species coded as andromonoecious, with a continuum from weakly andromonoecious species with a low proportion of staminate flowers to strongly andromonoecious species with a large number of staminate flowers and a single hermaphroditic flower that sets fruit (Whalen & Costich, 1986; Miller & Diggle, 2007). Future studies in large clades where andromonoecy is common but not predominant (e.g., EHS clade) will be useful in refining our knowledge of sexual system transitions in *Solanum*; both additional taxa and sequences will be needed to untangle relationships in the species-rich EHS clade where more detailed relationships are not yet clear. Small clades exhibiting the entire range of sexual systems (e.g., the Australian groups under study by Martine & al., 2019) could be used to understand the role of flower morphology and pollinator dynamics in sexual system evolution in relation to ecological conditions (e.g., Quesada-Aguilar & al., 2008).

### Future research

xxx

## Conclusion

Our analyses provide a systematic review and synthesis of morphological diversity in the large and economically important genus *Solanum* at clade level using the most up-to-date and well-sampled phylogeny, thus expanding on previous works both in terms of taxa and traits (see Knapp & al. 1998; Knapp, 2001, 2002b, 2002c; Bohs & al. 2007). Traits related to plant structure (growth form and sympodial unit structure), plant defence (glandular trichomes), pollination (corolla shape and colour), and dispersal (fruit colour) are identified as the most evolutionarily labile traits in *Solanum*. Ten further traits relating to life form (specialised underground organs), defence (prickle type, trichome structure), leaf division, inflorescence position and branching, sexual system, stamen hetermorphism and fruit morphology show signal of evolutionary lability with 50-100 transitions observed across *Solanum*. Nine traits are more conserved and can be used to assemble combinations of traits that define informally named clades in the genus. Morphological definition of infrageneric clades in the mega-diverse and globally distributed *Solanum* will provide help for species-level identification as well as enable us to confidently predict phylogenetic placement of unsampled species.

Our overview provides a ‘backbone’ for studying morphological evolution in *Solanum* and for designing future experiments. Data from molecular genetics will enable detailed understanding of the evolution of pathways involved in each trait (Smith & al., 2020) and is still largely lacking for most traits. We emphasise, however, that many of the broad categories used in our study are oversimplifications of the continuous variation observed in *Solanum*, and confirmation of character homology will be needed in many places. This will include observations of developmental sequences from species across the clades undertaking genus-wide investigation of the genetic/metabolic pathways of several key traits (e.g., leaf division, corolla and stamen bilateral symmetry, corolla colour, sexual systems). Studying morphological diversity in such a large and morphologically variable group as *Solanum* is challenging, and we hope that our overview provides a guide for next steps and a way to further target species for future study. Our results are limited by species sampling and resolution of our phylogeny; the number and position of the trait shifts will likely increase as sampling and resolution of the underlying phylogeny increases. Increased sampling is especially needed especially in the most diverse clades of *Solanum* (e.g., Brevantherum, Dulcamaroid, Geminata, Torva, and EHS clades) both in terms of species and genes to further refine and better understand phylogeny and evolution of this globally important genus.

## Supporting information

SI Table 1 List of Solanum clade

SI Table 2 Morphological data matrix

SI Table 3 Results of ancestral state reconstruction

SI Table 4 Solanum species per clade

## Acknowledgements

We thank National Science Foundation (NSF) grant (to LB & SK) ‘PBI *Solanum* – a world treatment’ DEB-0316614, Sibbald Trust Fellowship (to RH), the Fundación Ceiba and the GEME lab (Agreement 566 from 2014 between the Universidad Nacional de Colombia (https://unal.edu.co/) and Colciencias (now called Minciencias https://minciencias.gov.co/) (to AO), and CNPq (427198/2016-0 and 422191/2021-3) and FAPESPA/CAPES (Proc. AUXPE 88881.159124/2017-01 to LG) for funding and supported the research, Rocío Deanna for discussions about character states, and João Renato Stehmann, Richard Olmstead, and Stacey Smith for critical yet insightful comments that helped to enhance the manuscript. We thank the following people for images: Mario Vallejo-Marín, Martin Gardner, David Rae, Paulina Hechenleitner, Pedro Acevedo Rodríguez, Maria S. Vorontsova, Rocío Deanna, and Michael Benedito.

**SI Table 1.** List of informally recognised clades of *Solanum*, including species-level sampling and branch support in the supermatrix phylogeny of Gagnon & al. (2022) shown for each clade. Numbers of currently known species expected to belong to each clade based on morphological affinity are shown, and details can be found for the full list of accepted species in SI Table 4. The most recent and relevant taxonomic treatment with species descriptions and identification keys are indicated, as well as all more detailed published molecular phylogenetic studies. See main text and figures for full description of clades and clade names. BS = bootstrap support; PP = posterior probability support.

**SI Table 2.** Morphological data matrix used in the ancestral state reconstruction analyses, including sources used for scoring states for each species under all traits.

**SI Table 3.** Details of all ancestral state reconstruction results for all 25 traits based on maximum likelihood estimate of stochastic mapping in SIMMAP with 200 simulations and randomisations, including all estimated transition rates of the underlying Markov models of morphological evolution of each trait.

**SI Table 4.** List of accepted *Solanum* species and their position within the informally described major and minor clades. The position is based on the molecular phylogeny by Gagnon & al. (2022) for species sampled thus far; where sequence data is still lacking, species have been placed in clades based on morphology.

