## Supplementary material for "Morphological trait evolution in *Solanum* (Solanaceae): evolutionary lability of key taxonomic characters": SI Table 1 List of Solanum clade

**SI Table 1.** List of *Solanum* clades, including species-level sampling and branch support in the supermatrix phylogeny of Gagnon & al. (2022) shown for each clade. Numbers of currently known species expected to belong to each clade based on morphological affinity are shown, and details can be found for the full list of accepted species in SI Table 1. The most recent and relevant taxonomic treatment with species descriptions and identification keys are indicated, as well as all more detailed published molecular phylogenetic studies. See main text and figures for full description of clades and clade names. BS = bootstrap support; PP = posterior probability support.

| **Main group** | **Major clade** | **Minor clade** | **Phylogenetic sampling/known species (%)** | **Branch support** | **Detailed taxonomic and phylogenetic studies** |
| --- | --- | --- | --- | --- | --- |
| Thelopodium | Thelopodium | Thelopodium | 3/3 (100) | BS 100, PP 1 | Knapp, 2000 |
| Grade I | Regmandra | Regmandra | 6/12 (50) | BS 100, PP 1 | Bennett, 2008 |
|  | VANAns | Valdiviense | 2/2 (100) | BS 100, PP 1 | Knapp, 2013 |
|  |  | African non-spiny (Ans) | 5/14 (36) | BS 54, PP 0.76 | Knapp & Vorontsova, 2016 |
|  |  | Normania | 2/3 (67) | BS 100, PP 1 | Francisco-Ortega & al., 1993; Bohs & Olmstead, 2001 |
|  |  | Archaesolanum | 8/8 (100) | BS 99, PP 1 | Symon, 1981, 1994; Poczai & al., 2011 |
|  | DulMo | Dulcamaroid | 25/43 (58) | BS 99, PP 1 | Knapp, 2013 |
|  |  | Morelloid | 65/79 (82) | BS 100, PP 1 | Barboza 2013; Särkinen & Knapp, 2016; Knapp & Särkinen, 2018; Knapp & al., 2019, 2020; Särkinen & al., 2015a, b, c, 2018 |
|  | Potato | Herpystichum | 10/10 (100) | BS 50, PP 0.25 | Tepe & Bohs, 2011 |
|  |  | Pteroidea | 10/10 (100) | BS 100, PP 1 | Knapp & Helgason, 1997; Tepe & Bohs, 2010; Tepe & al., 2016 |
|  |  | *S. oxycoccoides* | 1/1(100) | BS 89, PP 1 | Bitter, 1919; Tepe & al., 2016 |
|  |  | Articulatum | 2/2 (100) | BS 100, PP 1 | Correll, 1962; Tepe & al., 2016 |
|  |  | Basarthrum | 10/16 (56) | BS 100, PP 1 | Correll, 1962; Seithe & Anderson, 1982; Anderson & Bernardello, 1991; Anderson & al., 2006a; Tepe & al., 2016 |
|  |  | Anarrhichomenum | 9/12 (75) | BS 100, PP 1 | Correll, 1962; Anderson & al., 1999; Tepe & al., 2012 |
|  |  | Etuberosum | 2/3 (67) | BS 100, PP 1 | Contreras & Spooner, 1999; Spooner & al., 2016 |
|  |  | Tomato | 14/17 (82) | BS 100, PP 1 | Peralta & al., 2008 |
|  |  | Petota | 61/113 (54) | BS 86, PP 1 | Spooner & al., 2004, 2008, 2016, 2019; Tepe & al., 2016 |
| Clade II | *S. anomalostemon* | *S. anomalostemon* | 1/1 (100) | BS 96, PP 1 | Knapp & Nee, 2009 |
|  | Brevantherum | Trachytrichium | 2/2 (100) | BS 100, PP 1 | Giacomin 2015 |
|  |  | Inornatum | 2/5 (40) | BS 100, PP 1 | Giacomin & Stehmann, 2014; Giacomin 2015 |
|  |  | Gonatotrichum | 7/7 (100) | BS 100, PP 1 | Giacomin & Stehmann, 2011; Giacomin 2015; Stern & Bohs, 2012; Stern & al., 2013 |
|  |  | Brevantherum | 29/80 (36) | BS 94, PP 1 | Roe, 1967, 1972; Freire de Carvalho & Machado, 1991; Giacomin 2015; Tovar & al., 2021 |
|  | Geminata | Reductum | 2/2 (100) | BS 100, PP 1 | Morton, 1976 |
|  |  | Geminata | 68/150 (45) | BS 97, PP 0.98 | Knapp, 2002, 2008; Knapp & al., 2015 |
|  | Cyphomandra | *S. graveolens* | 1/1 (100) | BS 100, PP 1 | Bohs, 1994 |
|  |  | Cyphomandropsis | 6/11 (55) | BS 40, PP - | Bohs, 2001, 2007; Falcão & Stehmann, 2018 |
|  |  | Pachyphylla | 33/42 (79) | BS 86, PP - | Bitter, 1913; Bohs, 1994, 1995, 2007; Bohs & Nelson, 1997; Stern & Bohs, 2009 |
|  | Wendlandii-Allophyllum | Allophyllum | 4/4 (100) | BS 32, PP - | Bohs, 1989, 1990; Nee & al., 2006 |
|  |  | Wendlandii | 5/9 (56) | BS 100, PP 1 | Clark & al., 2015; Cuevas-Guzmán & Núñez-López, 2015 |
|  | Nemorense | Nemorense | 4/4 (100) | BS 92, PP 1 | Child, 1983 |
|  | Leptostemonum | *S. polygamum* | 1/1 (100) | BS 96, PP 1 | Vahl, 1794; Anderson & al., 2015 |
|  |  | Lasiocarpa | 12/12 (100) | BS 100, PP 1 | Whalen & al., 1981; Bohs, 2004; Miller & Diggle, 2007; Aubriot & al., 2022 |
|  |  | Acanthophora | 13/22 (59) | BS 100, PP 1 | Nee, 1986, 1991; Levin & al., 2005, 2006; Miller & Diggle, 2007; Stern & al., 2011; Chiarini & Mentz, 2012 |
|  |  | Gardneri | 8/10 (80) | BS 100, PP 0.99 | Levin & al., 2006; Stern & al., 2011; Silva Sampaio & al., 2021 |
|  |  | Thomasiifolium | 4/9 (44) | BS 86, PP 0.94 | Levin & al., 2006; Stern & al., 2011 |
|  |  | Erythrotrichum | 14/35 (40) | BS 98, PP 0.57 | Agra, 2004, 2008 |
|  |  | Sisymbriifolium | 2/2 (100) | BS 100, PP 1 | Levin & al., 2006; Stern & al., 2011 |
|  |  | Crinitum | 10/23 (43) | BS 94, PP 0.98 | Roe, 1966; Levin & al., 2006; Agra & Stehmann, 2016; Gouvêa & al., 2019 |
|  |  | Androceras | 15/16 (94) | BS 100, PP 0.67 | Whalen, 1979, 1986; Stern & al., 2010, 2014 |
|  |  | *S. campechiense* | 1/1 (100) | BS 96, PP 0.29 | Levin & al., 2006; Stern & al., 2011 |
|  |  | Carolinense | 10/14 (71) | BS 96, PP 1 | Levin & al., 2006; Stern & al., 2011; Wahlert & al., 2014, 2015 |
|  |  | Bahamense | 3/3 (100) | BS 89, PP 0.99 | Levin & al., 2006; Stern & al., 2011;  Strickland-Constable & al. 2010 |
|  |  | Micracantha | 9/14 (64) | BS 100, PP 1 | Levin & al., 2006; Stern & al., 2011 |
|  |  | Asterophorum | 2/4 (50) | BS 100, PP 1 | Gouvêa & Stehmann, 2019 |
|  |  | *S. multispinum* | 1/1 (100) | BS 74, PP 1 | Levin & al., 2006; Stern & al., 2011 |
|  |  | Torva | 34/55 (62) | BS 97, PP 1 | Levin & al., 2006; Stern, 2014; Stern & al., 2011; Aubriot & al., 2022 |
|  |  | *S. euacanthum* | 1/1 (100) | BS 97, PP 1 | Barboza, 2013 |
|  |  | Elaeagnifolium | 5/5 (100) | BS 100, PP 1 | Knapp & al., 2017 |
|  |  | Eastern Hemisphere Spiny (EHS) | 198/336 (59) | BS 95, PP 1 | Whalen, 1984; Symon, 1981, 1985; Bean, 2001, 2004, 2011, 2012, 2014, 2016a, b; Anderson & al., 2006a,b; Levin & al., 2006; Martine & al., 2006, 2009, 2016, 2019; Vorontsova & al., 2010, 2013; Stern & al., 2011; Knapp & al., 2013; Aubriot & al., 2022; Vorontsova & Knapp, 2016; McDonnell & al., 2019 |
